# Assembly-Activating Protein Phase Separation Properties Are Required for Adeno-Associated Virus Type 2 Assembly

**DOI:** 10.1101/2025.03.10.642135

**Authors:** Janine Vetter, Manuel Kley, Catherine Eichwald, Cornel Fraefel

## Abstract

Adeno-associated virus serotype 2 (AAV2), a non-pathogenic parvovirus reliant on helper viruses, is studied extensively as a potential gene delivery vector. A +1 open reading frame within the cap gene encodes a nonstructural protein of 204-amino-acids termed assembly-activating protein (AAP), which has been attributed a critical role in transporting the viral capsid protein VP3 into the nucleolus for assembly. However, AAP remains poorly characterized because of its relatively late discovery and lack of commercial antibodies. In the absence of other virus proteins, AAP localizes in the nucleolus due to five redundant nuclear and nucleolar localization signals. Additionally, AAP, a predicted intrinsically disordered protein, forms spontaneous dose-dependent nuclear globular condensates, a trait of liquid-liquid phase separated inclusions. Consistent with LLPS biophysical properties, the AAP condensates recovered rapidly from photobleaching and are sensitive to aliphatic diol treatment—moreover, AAP self-oligomerizes. We produced an AAP-specific antibody to analyze the role of this protein during productive AAV2 replication. In this context, we observed that AAP also forms nuclear globular condensates with LLPS biophysical properties in cells co-infected with AAV2 and either herpes simplex virus type 1 (HSV-1) or adenovirus type 5 (AdV-5) as the helper viruses. The screening of AAP deletion mutants revealed that the N-terminal region (amino acids 1-61) is necessary for condensate formation and self-oligomerization. Interestingly, this AAP region contains a predicted alpha-helix spanning amino acids 16 to 45. The substitution in this region of the hydrophobic residues by alanines drastically impaired AAP-LLPS biophysical properties and its ability to trigger AAV2 capsid assembly. Identifying the amino acids involved in assembly and LLPS may improve AAV vector production.

**Author Summary:** Adeno-associated virus serotype 2 (AAV2) is a non-pathogenic virus extensively studied for its potential in gene therapy. It relies on a protein called assembly-activating protein (AAP) to transport its capsid protein, VP3, to the nucleolus for assembly. The 204-amino-acid AAP is not well characterized because it was discovered only relatively recently and commercial antibodies are not availabe, making it challenging to study. Here, we demonstrate that AAP localizes in the nucleolus and forms globular condensates through liquid-liquid phase separation (LLPS), a property characterized by rapid recovery from photobleaching and sensitivity to aliphatic diol treatment. Additionally, we prepared a specific antibody to study AAP during AAV2 co-infection with helper viruses like herpes simplex virus type 1 (HSV-1) or adenovirus type 5 (AdV-5). We found that AAP also forms nuclear condensates with LLPS properties in co-infected cells. We demonstrate that the N-terminal region of AAP (amino acids 1-61) is crucial for condensate formation and self-oligomerization. Within this region, a predicted alpha-helix (amino acids 16-45) is essential, as substituting its hydrophobic residues with alanines significantly impaired the LLPS properties of AAP and its ability to facilitate AAV2 capsid assembly. Identifying these key amino acids may enhance AAV vector production for gene therapy applications.

## Introduction

Adeno-associated virus (AAV) is a small, non-pathogenic parvovirus widely studied as a delivery vehicle for gene therapy [1]. AAV relies on a helper virus for productive replication [2]. When not aided by a helper virus, AAV can integrate its genome into a preferred site on human chromosome 19, termed the adeno-associated virus pre-integration site 1 (AAVS1), or it can persist in the nucleus in an episomal form [3–5]. However, upon co-infection with a helper virus such as herpes simplex virus type 1 (HSV-1), adenovirus type 5 (AdV-5), or human papillomavirus type 16, AAV initiates a lytic replication cycle, resulting in the generation of virus progeny [2]. Throughout the infection process, AAV is known to induce the formation of different nuclear compartments in infected cells [6].

The genome of AAV is a single-stranded DNA of about 4.7 kilobases comprising two large open reading frames (ORFs), flanked by 145 nucleotides long inverted terminal repeats (ITRs) on both ends [7, 8]. The two ORFs encode the nonstructural and structural proteins involved in various functions. The first ORF is driven by two promoters (p5 and p19) and contains one site for alternative splicing, leading to the expression of four nonstructural proteins: Rep78, Rep52, Rep68, and Rep40 [9]. Rep78 and Rep68 are involved in AAV replication, while Rep52 and Rep40 are crucial in genome packaging [2].

The second ORF, containing the cap gene, is driven by one promoter (p40) and encodes three in-frame proteins, VP1, VP2, and VP3, initiated from three different start codons [10]. The icosahedral capsid of AAV is composed of 60 viral proteins arranged into an icosahedral structure (T=1), with VP1, VP2, and VP3 present in a 1:1:10 molar ratio [11]. The cap gene also encodes two frame-shifted proteins, the membrane-associated accessory protein (MAAP) [12] and the assembly-activating protein (AAP) [13, 14].

AAP, encoded from a +1 open reading frame, is 204 amino acids long and plays a critical role in capsid assembly [14]. The assembled AAV capsids were found to co-localize with AAP in different nuclear locations, such as the nucleoplasm, nucleolus, or clustered around the nuclear membrane, depending on the AAV serotype [15]. AAP from AAV serotype 2 (AAP2) is responsible for transporting VP3 into the nucleolus. However, simply targeting VP3 to the nucleolus by adding nucleolar localization signals is insufficient to induce the assembly of virus-like particles [14]. This indicates additional crucial roles of AAP, including stabilization of the VP proteins [16]. AAP remains poorly characterized because of its relatively late discovery and lack of commercial antibodies. AAV serotype 2 (AAV2) assembly significantly relies on AAP, whereas the 13 naturally occurring serotypes exhibit varying levels of dependence. Serotypes 4, 5, and 11 do not require AAP for assembly [15, 17]. The expression levels of AAP2 increase steadily during AAV2 infection in the presence of adenovirus helper plasmids, followed by an abrupt drop after 24 h post-infection [12].

AAP comprises three main regions: the N-terminus, the central domain, and the C-terminus. The N-terminus contains a hydrophobic region and a conserved core, which seem essential for the assembly function [16, 18]. Furthermore, the central region of AAP is rich in threonines and serines, suggesting a high potential for phosphorylation. The C-terminus contains five redundant nuclear and nucleolar localization signals [19].

Recently, numerous studies have revealed that the specialized membrane-less compartments formed during viral infection are predominantly liquid-liquid phase separated (LLPS) condensates [20–26]. Membrane-less condensates can provide optimal concentrations of viral and cellular factors for virus replication and simultaneously shield from cellular innate immune responses [27]. LLPS is a physical process in which two or more substances separate into distinct phases due to energetically favorable interactions [28]. Proteins undergoing phase separation often exhibit several of these characteristics: an intrinsically disordered region (IDR), a tendency to self-oligomerize due to multivalency, an RNA-binding site, or high phosphorylation levels [28–32]. Important hallmarks of LLPS include the formation of globular structures, coalescence events, and rapid recovery from photobleaching (FRAP) [28, 33]. In this study, we demonstrate that AAP2 confers liquid-like properties to viral compartments and that LLPS and AAV2 capsid assembly depend on AAP2 self-oligomerization, conferred by hydrophobic amino acids in the N-terminus.

## Results

### AAP2 shows the hallmarks of an LLPS protein

It is well known that AAV2 relies on the presence of AAP2 to successfully assemble its capsid, with AAP2 forming compartments in the nucleus [14]. Many viral compartments are biomolecular condensates (BCs) with LLPS properties due to large IDRs that drive their assembly and maintenance [28, 29]. We investigated whether AAP2 inclusions exhibit behavior consistent with BCs by studying their LLPS properties. As a first step, we performed an *in-silico* analysis using the VSL2 algorithm from the Predictor of Natural Disordered Regions (PONDR) website. Consistent with AAP2 compartments forming by phase separation, large parts of the AAP2 core and C-terminal regions are predicted to be disordered (81.86% of the protein) (**Fig 1A**). Then, we expressed different versions of fluorescently labeled AAP2 in BJ cells, either fused to a green fluorescent protein (GFP) or monomeric kusabira orange (mKO) and analyzed their subcellular distribution. Confocal laser scanning microscopy (CLSM) followed by three-dimensional reconstruction revealed the formation of AAP2-GFP and AA2P-mKO globular condensates in the nucleus (**Fig 1B**). It appears that two types of inclusions can form: larger ones that co-localize with nucleolar proteins such as nucleophosmin and fibrillarin and smaller ones that are dispersed in the nucleoplasm and co-localize with nucleophosmin but not fibrillarin (**Fig S1, A and B**). Additionally, the co-expression of AAP2-GFP and AAP2-mKO leads to the formation of mixed compartments (**Fig 1B**). Of note, tagging AAP2 at the N-terminus or C-terminus did not impair its ability to form condensates in the nucleus (**Fig S1C**). We calculated mean solidity values (area/convex area) of 0.8 and 0.0 in cells transfected with AAP2-GFP and GFP, respectively (**Fig 1C**). The solidity value is a shape descriptor calculated by dividing the area of a shape by its convex area. Therefore, a solidity value approaching one indicates a round BC, pointing to surface tension in the compartment and phase separation.

**Fig 1.**
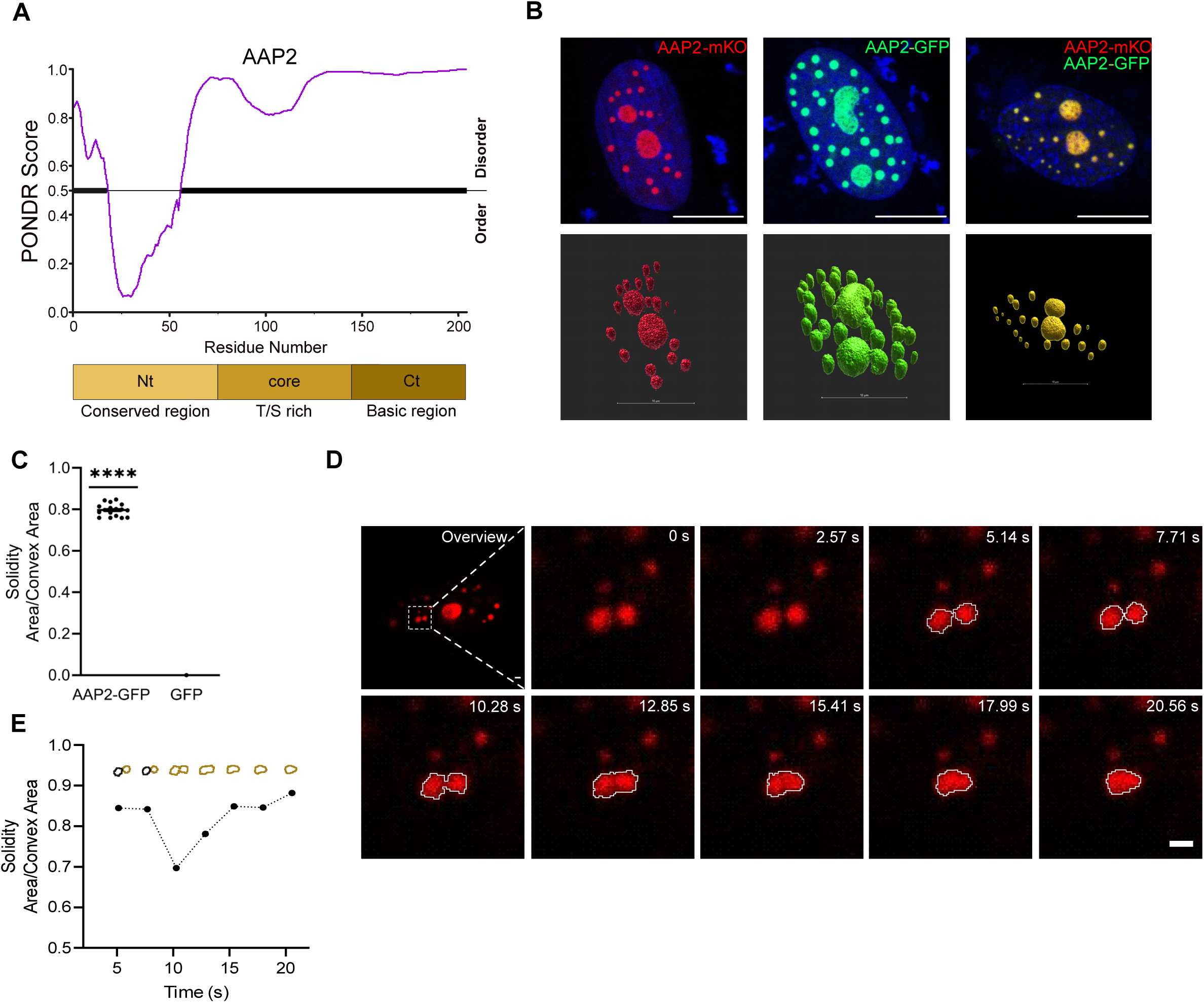
AAP2 forms nuclear globular condensates. (A) Plot for IDR prediction of AAP2. The bold part of the grey line highlights the disordered region of the protein. The main AAP2 regions are shown below. (B) Fluorescence images of BJ cells expressing AAP2-mKO (red, left column), AAP2-GFP (green, middle column), and both (yellow merge, right column). At 16 hpt, the cells were fixed, and nuclei were stained with DAPI (blue). The top and bottom rows correspond to a single focal plane and three-dimensional reconstruction. The scale bar is 10 μm. (C) Plot for the solidity ratios of AAP2-GFP condensates at 16 hpt. A solidity ratio close to 1 indicates a round shape. The data represent the mean ± SD; One-sample t-test with a hypothetical mean value of 0, (****) p < 0.0001, n ≥ 17. (D) Time-lapse images of BJ cells expressing AAP2-mKO (red). The samples were acquired at 16 hpt with intervals of 2.57 sec. The dashed open box in the first panel corresponds to the enlarged images shown in the subsequent panels. The scale bar is 1 μm. (E) Plot for solidity ratios of the fusing AAP-mKO droplets from panel D. The ochre-shaped droplets correspond to white lines in panel D.

Furthermore, we observed fusion events of AAP2-mKO condensates in live-cell microscopy (**Fig 1D**), another hallmark of phase separation [33]. The condensates appeared mobile and, upon contact, rapidly coalesced and relaxed back into spheres with a high solidity ratio **(Fig 1E)**, similar to other LLPS BCs [23, 34, 35]. In summary, AAP2 has large, disordered regions and forms miscible globular inclusions that can coalesce, exhibiting LLPS properties.

### AAP2 condensates are dynamic

BCs can be characterized by their sensitivity to 1,6-hexanediol (1,6-HD), an aliphatic diol dissolving weak molecular bonds in LLPS condensates [24, 36–41]. To quantitatively characterize the properties of AAP2 condensates, BJ cells expressing AAP2-GFP were treated with 1,6-HD. As expected, a 5-minute treatment with 1,6-HD led to the partial dissolution of AAP2 condensates (**Fig 2A**), denoted by a significant decrease in the area (**Fig 2B**) and solidity (**Fig 2C**). Furthermore, we performed fluorescence recovery after photobleaching (FRAP) experiments (**Fig 2, D-H**) to characterize the LLPS properties of the AAP2 condensates. Condensates with liquid-like properties are expected to rapidly recover after the photobleaching, while this is prevented in solid compartments due to a lack of mobility [42]. We bleached three different size categories of condensates (< 1.5 µm, = 1.5 µm, > 1.5 µm) with a bleaching diameter of 1.5 µm (**Fig 2E)**, allowing us to visualize simultaneously *i)* the rates of diffusion within the condensate and *ii)* the turnover rate between the condensates and their nuclear environment. We observed rapid recovery in all three condensate size categories (within 40-60s) (**Fig 2 F-H**) but no difference between the mobile fraction of the condensates (**Fig 2H**), suggesting that the same number of molecules are mobile in all condensates. Interestingly, the half-maximal recovery time of the large condensates was significantly longer than that of the smaller condensates (**Fig 2G**). This outcome suggests a different diffusion rate within the BC compared to the turnover rate between condensates and their nuclear environment, probably due to differences in material properties of the two environments.

**Fig 2.**
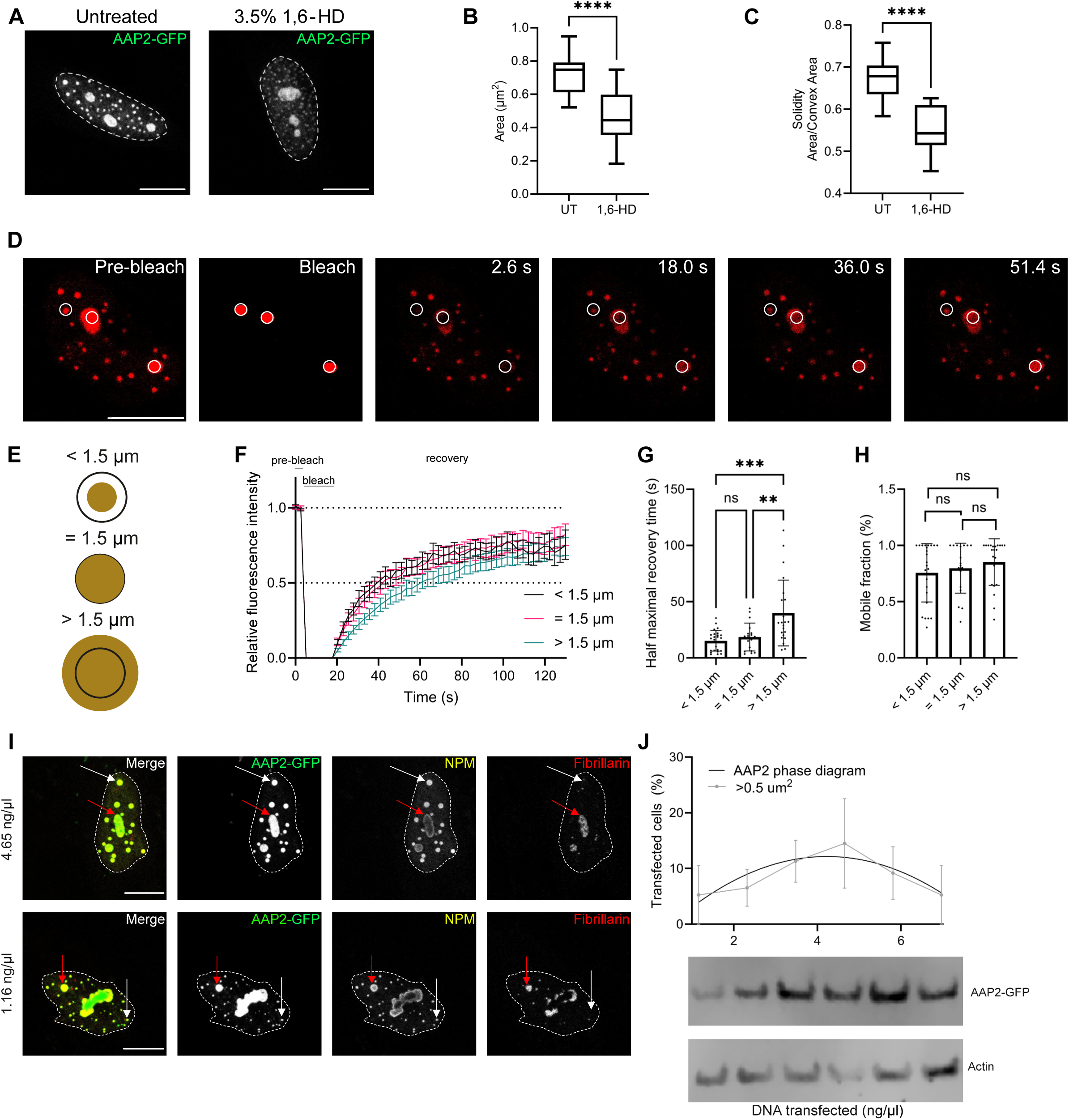
AAP2 condensates show hallmarks of LLPS. (A) Representative immunofluorescence images of BJ cells expressing AAP2-GFP untreated (left image) or treated (right image) with1,6-HD. At 16 hpt, the cells were untreated and treated with 3.5% 1,6-HD for 5 min, fixed with 2% paraformaldehyde, and nuclei stained with DAPI (white dashed line). The scale bar is 10 μm. Plot for size (B) and solidity ratio (C) of AAP2 condensates per nucleus untreated (UT) and treated with 1,6-HD. The data correspond to the mean ± SD, Welch’s t-test, (****) p < 0.0001, n≥15. (D) Representative time-lapse fluorescence images of FRAP measurements from BJ cells expressing AAP2-mKO (red). At 16 hpt, the AAP2-mKO condensates were in the indicated areas at pre-bleach, bleached, and recovery time conditions (white circles), followed by monitoring for 51.4 s. The images were acquired at 2.6 s intervals. The scale bar is 10 μm. (E) Schematic representation of sorted bleached areas in AAP2-mKO condensates. The AAP2-mKO condensates (ochre) were sorted as either small (<1.5 µm, top), medium (= 1.5 µm), or large (>1.5 µm). The bleached area had a constant radius of 1.5 μm (black circle). (F) FRAP recovery curves of single AAP2-mKO condensates measuring <1.5 (grey), =1.5 (red), and >1.5 (blue) µm. The data represent the mean ± SEM. n ≥ 18 for each measurement. The dotted lines indicate the recovery time until 50% and 100% relative fluorescence intensity. Plots indicating the mobile fraction (G) and recovery half-time (H) means of the different-sized AAP2-mKO condensates after photobleaching. The data represent the mean values ± SD; one-way ANOVA, (**)p<0.01 and (***) p<0.001. (I) Immunofluorescence images of BJ cells expressing AAP2-GFP (green) transfected at increasing concentrations of pCI-AAP2-GFP DNA. At 16 hpt, the cells were fixed, followed by immunostaining for the detection of nucleolar proteins nucleophosmin (anti-NPM, yellow) and fibrillarin (anti-fibrillarin, red). The nuclei were stained with DAPI (white dashed line). The red and white arrows point to the nucleolar and extranucleolar AAP2-GFP condensates, respectively. The scale bar is 10 μm. (J) Plot for the phase diagram of DNA transfected cells (ng/µl) in (I). The grey line corresponds to the fraction of extranucleolar condensates larger than 0.5 µm^2^. The data corresponds to three independent experiments. At the bottom, immunoblotting is performed to examine the expression of AAP2-GFP (anti-GFP). β-actin was used as a loading control.

The proteins driving LLPS can display this behavior in a concentration-dependent manner [24, 43]. This hallmark of phase separation was observed when we transfected BJ cells with low (1.16 ng/µl) or high (4.65 ng/µl) DNA concentrations of AAP2-GFP plasmid. At 16 h post-transfection, the cells were stained for detection of nucleophosmin (NPM, yellow) and fibrillarin (red) to distinguish between large nucleolar and smaller extranucleolar AAP2 condensates, respectively (**Fig 2I**). No difference in nucleolar AAP2 BCs was observed, as denoted by the colocalization of NPM and fibrillarin signals, regardless of the DNA concentration. However, the extranucleolar AAP2 BCs, as denoted by the lack of fibrillarin signal, demonstrate a notable increase in size as the DNA concentration increases. Moreover, BJ cells were transfected with increasing amounts of AAP2-GFP DNA, and the percentage of extranucleolar AAP2-GFP condensates larger than 0.5 µm^2^ was determined (**Fig 2J**). As illustrated in a phase diagram denoting the concentration dependence of BCs, the percentage of transfected cells varied depending on the amount of DNA used, with maximum phase separation at a DNA concentration of 4.23 ng/µl (**Fig S1D**). These experiments suggest that the expression of AAP2 can induce the formation of phase-separated condensates.

### AAP2 condensates forming upon infection with AAV2 show hallmarks of LLPS

After confirming that the overexpression of AAP2 leads to LLPS BC formation, we aimed to investigate their occurrence in cells co-infected with AAV2 and a helper virus. For this purpose, we generated an antibody targeting the AAP2 C-terminus region from amino acids 179 to 194, which specifically recognizes AAP2 in both immunofluorescence and immunoblotting (**Fig S2, A-B**). Of note, in immunofluorescence assays of cells co-infected with AAV2 and replication-competent HSV-1, the anti-AAP2 antibody presents unspecific cytoplasmic aggregates; however, the nuclear signal is specific for AAP2 (**Fig S2C**). Next, we compared and quantified AAP2 nuclear condensate formation in co-infected (AAV2 and either HSV-1 or AdV-5) Rep-positive cells at three time points, 16, 24, and 40 hpi (**Fig 3, A-F**). The distribution of Rep varies according to the helper virus used. In AdV-5 co-infected cells, the Rep staining surrounded the globular AAP2 inclusions (**Fig 3A**), while in HSV-1 co-infected cells, AAP2 was diffused in the nucleoplasm (**Fig 3B**). This specific Rep phenotype was not an artifact of AAP2 antibody staining, as the same Rep pattern was observed also upon staining with the Rep-specific antibody alone (**Fig S2D**). Upon AdV-5 co-infection, the numbers of AAP2 condensates were higher at 24 hpi than at 40 hpi (**Fig 3C**), but the solidity values were comparable at the two-time points (**Fig 3E**). In cells co-infected with AAV2 and HSV-1, AAP2 condensates were formed by 16 hpi. While their numbers did not increase by 24 hpi **(Fig 3D),** a decrease in inclusion solidity values was observed in association with changes in morphology **(Fig 3F).**

**Fig 3.**
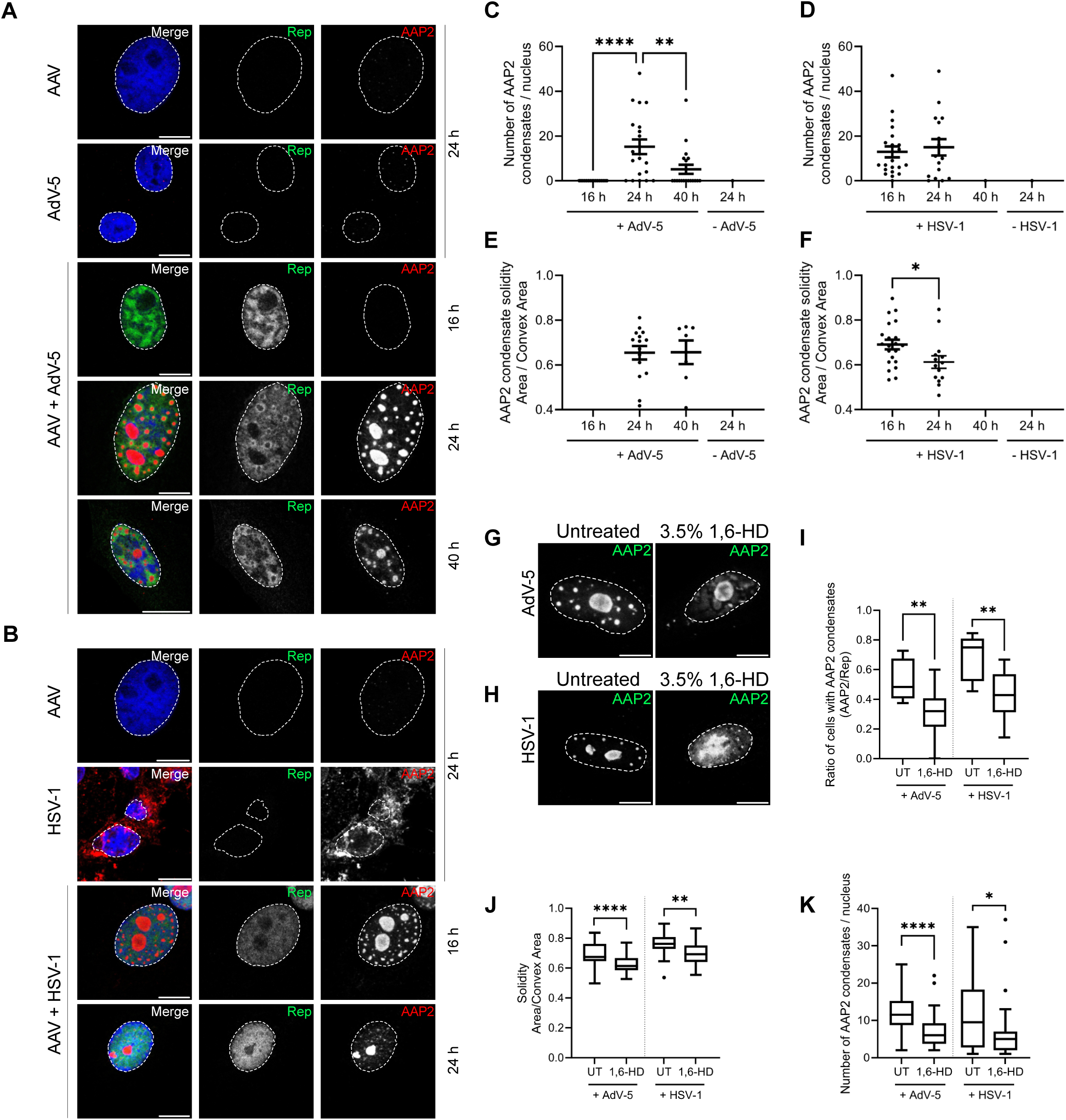
AAP2 condensates in infected cells show hallmarks of LLPS. Immunofluorescence images of Vero cells co-infected with AAV2 (MOI 500) and either AdV-5 (MOI 0.1) (A) or HSV-1 (MOI 0.1) (B). At 16, 24, and 40 hpi, the cells were paraformaldehyde-fixed, followed by immunostaining for the detection of Rep proteins (anti-Rep, green, middle columns) and AAP2 (anti-AAP, red, right columns). The nuclei were stained with DAPI (blue, dashed white line). Merged images are presented in the left column. The scale bar is 10 µm. Plot for the numbers and solidity of nuclear AAP2 condensates in cells co-infected with AAV2 and either AdV-5 (C, E) or HSV-1 (D, F). The data represent the mean values ± SEM; one-way ANOVA, (*) p< 0.05, (**) p< 0.01, (****) p < 0.0001, n ≥ 15. (G, H) Immunofluorescence images of Vero cells co-infected with AAV2 (MOI 500) and the indicated helper virus (MOI 0.1). At 24 hpi (AdV-5, G) or 16 hpi (HSV-1, (H), the cells were untreated (left image) or treated (right image) for 5 min with 3.5% 1,6-HD. Then, the cells were immediately fixed with 2% paraformaldehyde, followed by immunostaining to detect AAP2 (anti-AAP, white). The nuclei were stained with DAPI (white dashed line). The scale bar is 10 μm. Tukey plots for quantification of the ratio of cells presenting AAP2 condensates (I), the solidity of AAP2 condensates (J), and the number of nuclear AAP2 condensates (K). Student’s t-test, (*) p< 0.05, (**) p< 0.01, (****)p < 0.0001, n ≥ 30

LLPS of AAP2 condensates were further characterized by their sensitivity to 1,6-HD treatment in co-infected cells. AAP2 condensates visibly dissolved within the 5 min of treatment with 1,6-HD, independently of the helper virus used (**Fig 3G and H**). The quantification of the condensates after 1,6-HD treatment revealed a significant decrease in the ratio of AAP2 condensates per Rep expressing cells (**Fig 3I**), in the number of AAP2 condensates per nucleus (**Fig 3K**), and globular morphology, as denoted by a decrease in their solidity (**Fig 3J**).

These findings demonstrate the dynamic nature and LLPS properties of AAP2 inclusions upon AAV2 coinfection with a helper virus. Therefore, AAP2 inclusions may be considered BCs.

### The AAP2 N-terminal region is necessary for the formation of BCs

Next, we screened a series of AAP2 deletion mutants fused in-frame to GFP (**Fig 4A**) to identify the AAP2 region responsible for condensate formation and phase separation. For this, AAP2 was divided into three regions corresponding to the N-terminus (Nt, amino acids 1 to 61), core (amino acids 62 to 143), and C-terminus (Ct, amino acids 144 to 204) in different combinations. The correct expression of all tested AAP2-GFP deletion mutants was verified by immunoblotting (**Fig S3 A and E**). The AAP2-GFP deletion mutants were transfected into BJ cells and analyzed by immunofluorescence for their ability to form condensates (**Fig 4B**). Not surprisingly, all deletion mutants containing the C-terminal region localized in the nucleoli since this region has been recognized previously to contain five redundant nuclear and nucleolar localization signals [19]. As expected, mutants lacking the C-terminus were mainly found in the cytoplasm due to the absence of a nuclear localization signal. Interestingly, all deletion mutants harboring the AAP2 N-terminal region formed condensates in the cytoplasm (Nt and Nt-core) or the nucleus (Nt-Ct). Therefore, we tested the LLPS properties of the N-terminus of AAP2 by treating cells expressing AAP(Nt)-GFP with 1,6-HD. The AAP(Nt)-GFP cytosolic inclusions were sensitive to 1,6-HD, dissolving within the 5 min of treatment (**Fig 4C**) and forming small and amorphous aggregates (**Fig 4D and E**). Moreover, AAP(Nt-core)-GFP globular inclusions formed in a dose-dependent manner as denoted in the phase diagram plot following cell transfection with increasing concentrations of plasmid DNA (**Fig S1 E and F**). Thus, the AAP2 N-terminal region from amino acids 1 to 60 is necessary for forming inclusions with LLPS properties.

**Fig 4.**
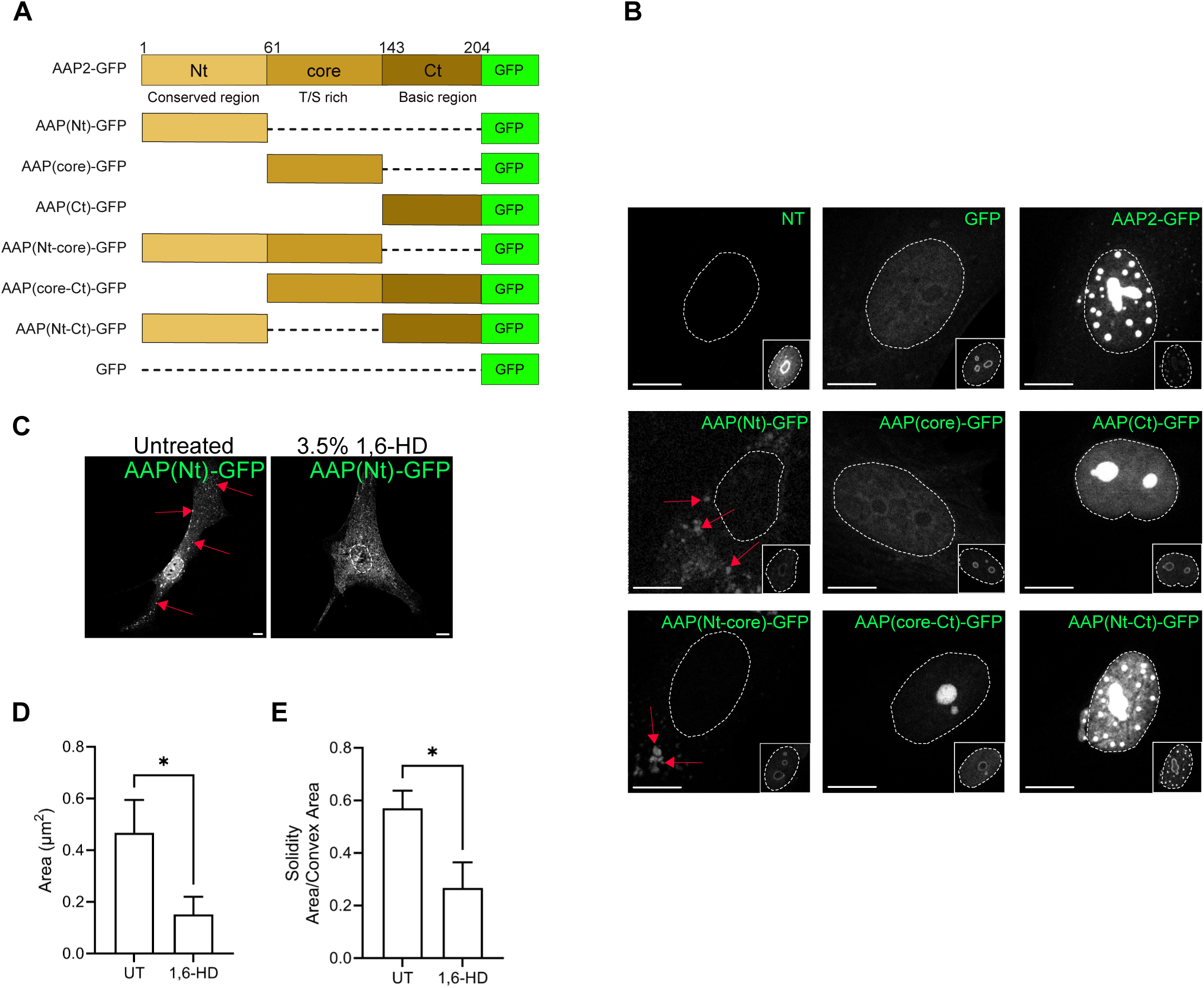
AAP(Nt)-GFP condensates show hallmarks of LLPS. (A) Schematic representation of the AAP2 deletion mutants fused to GFP. AAP2 was divided into three main regions: Nt (amino acids 1 to 61), core (amino acids 62 to 143), and Ct (amino acids 144 to 204). Nt, core, and Ct are conserved, T/S rich and basic regions, respectively. (B) Immunofluorescence images of BJ cells expressing the indicated AAP2-GFP deletion mutants (white). At 16 hpt, the cells were fixed, followed by immunostaining to detect nucleophosmin (anti-NPM, white, insets right bottom corner). The nuclei were stained with DAPI (white dashed line). The red arrows point to extranuclear AAP2-GFP condensates. The scale bar is 10 μm. (C) Immunofluorescence images of BJ cells expressing AAP(Nt)-GFP untreated (left image) and treated (right image) with 1,6-HD. At 16 hpt, the cells were untreated or treated for 5 min with 3.5% 1,6-HD. The cells were immediately fixed, and the nuclei were stained with DAPI (white dashed line). The red arrows point to extranuclear AAP(Nt)-GFP condensates. The scale bar is 10 μm. Plots for the quantification of AAP(Nt)-GFP condensates size (D) and solidity ratio (E). The data represent the mean ± SEM; Student’s t-test, (*) p < 0.05, n ≥13.

### AAP2 can self-oligomerize

Self-oligomerization is a trait of proteins driving LLPS in BCs [44]. It has been demonstrated that AAP2 forms high-molecular-weight oligomers; however, it remains unclear whether these are self-oligomers [16]. Since the N-terminal region of AAP2 is predicted to be partially ordered (**Fig 1A**), we hypothesized that this AAP2 region is required for self-oligomerization. To address this, we used a reovirus-derived *in vivo* protein-protein interaction platform to evaluate AAP2 self-oligomerization [45, 46]. This interaction assay employs the C-terminal region (amino acids 471 to 721) of the reovirus protein µNS, which forms cytosolic inclusions. Fusing the C-terminus of µNS to a protein of interest serves as bait for assessing if a positive protein-protein interaction occurs. The positive interaction can be visualized directly under a fluorescence microscope. In this context, AAP2-GFP localizes in the nucleus, forming inclusions when expressed alone or in co-expression with mCherry-µNS (**Fig 5A**). However, the co-expression of AAP2-GFP with AAP2-mCherry-µNS leads to cytosolic localization of AAP2-GFP in AAP2-mCherry-µNS platforms, thereby validating this assay to study the self-association of AAP2. Next, AAP2-GFP deletion mutants (**Fig 4A**) were tested for their ability to localize in AAP2-mCherry-µNS platforms. As expected, none of the tested AAP2-GFP deletion mutants localize with control mCherry-µNS platforms (**Fig S3D**). Interestingly, deletion mutants containing the N-terminal region of AAP2, such as AAP(Nt)-GFP or AAP(Nt-core)-GFP, co-localized with AAP2-mCherry-µNS platforms (**Fig 5B**). Meanwhile, deletion mutants lacking the N-terminal region of AAP2 resulted in a negative colocalization with AAP2-mCherry-µNS (**Fig 5B)**. To confirm the *in vivo* protein-protein interaction assay results, we developed a capture ELISA. In this assay, the well is coated with a polyclonal anti-GFP antibody followed by incubation with extracts of cells co-expressing AAP2-GFP, AAP(Nt)-GFP, or AAP(core-Ct)-GFP with AAP2-Flag. The association of AAP2-Flag with the AAP2 deletion mutants fused to GFP was subsequently detected with a monoclonal anti-Flag antibody. As anticipated, AAP2-GFP successfully interacts with AAP2-Flag. Furthermore, AAP(Nt)-GFP exhibited significantly higher self-oligomerization levels compared to AAP(core-Ct)-GFP **(Fig 5C)**. These combined results indicate the importance of the N-terminus region of AAP2 (amino acid 1 to 60) for AAP2 self-oligomerization.

**Fig 5.**
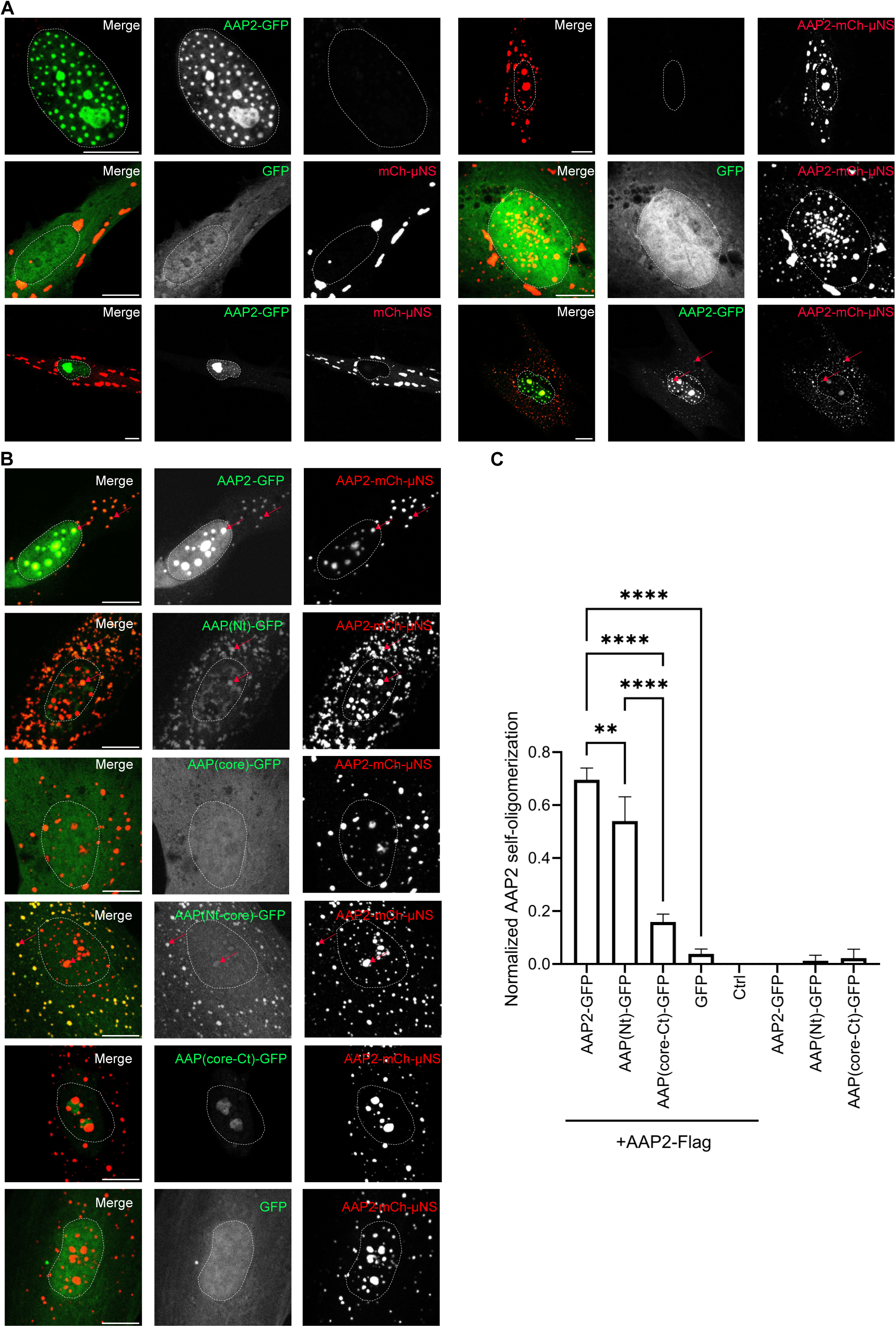
AAP(Nt)-GFP can self-oligomerize. In vivo protein-protein interaction using reovirus platform in which GFP tagged protein is “fish,” and mCherry-µNS tagged protein is “bait.” (A) At 16 hpt, BJ cells expressing AAP2-GFP (left top row) or AAP2-mCherry-µNS (right top row) or co-expressing either GFP (middle row) or AAP2-GFP (bottom row) with either mCherry-µNS (left panel) or AAP2-mCherry-µNS (right panel) were fixed, stained with DAPI (dashed white line) and analyzed by fluorescence microscopy. A merged image is shown in the left column. The red arrows point to the colocalization of AAP2-GFP in AAP2-mCherry-µNS platforms. The scale bar is 10 µm. (B) At 16 hpt, BJ cells expressing “fish” AAP2-GFP or indicated deletion mutants (green, middle column) with “bait” AAP2-mCherry-µNS (red, right column) were fixed, stained with DAPI (dashed white line) and analyzed at the fluorescence microscope. The red arrows point to the colocalization of AAP2-GFP or deletion mutants in AAP2-mCherry-µNS platforms. The scale bar is 10 µm. (C) Plot for the detection of the association of AAP2-GFP full-length or indicated deletion mutants to AAP2-Flag. The positive association was detected using a monoclonal anti-Flag antibody followed by incubation with anti-mouse conjugated to HRP. The data represents the mean ± SD of three independent experiments; one-way ANOVA, (****) p<0.0001, (**) p<0.01.

### Mutation of hydrophobic resides in the N-terminus of AAP2 to alanine prevents self-oligomerization, LLPS, and capsid assembly

Consistent with a PONDR prediction of a partially ordered N-terminal region of AAP2 (**Fig 1A**), AlphaFold2 analysis predicts an α-helix in residues 21 to 40 with a relatively high likelihood (**Fig S4A i**). This α-helix features hydrophobic amino acids exposed on one side. We hypothesized that these hydrophobic residues could be involved in AAP2 self-oligomerization. To address this, the bulky amino acids present in the predicted α-helix of AAP2 were divided into three regions, A, B, and C, and then mutated individually (A, B, C) or combined (AB, AC, BC, and PM) (**Fig 6A, Fig S4B and C**). Most of these mutants (**Fig S4D and E**) showed abrogation of extranucleolar condensate formation, except for mutants B and C, which did not lose the ability to induced BC formation. Consistent with this observation, AAP(PM)-GFP harboring point mutations in all three groups did not form BC, showing diffused signal in the nucleoplasm **(Fig 6B, first row).** This pattern matches the one for for AAP(core-Ct)-GFP **(Fig 4A and Fig 6B, second row)**, which also did not form extranucleolar BCs in BJ cells. Similarly, untagged AAP(PM), when detected with the specific anti-AAP antibody, did not form inclusions and was diffused in the nucleoplasm **(Fig 6B, second row).** Thus, AAP2 and AAP(PM) localization patterns are similar to APP2-GFP and AAP(PM)-GFP, respectively. Furthermore, also the quantification of the number of AAP(PM)-GFP nuclear condensates shows a significant decrease compared to the AAP2-GFP(**Fig 6C).** These results align with the inability of these point mutants to associate with wild-type AAP2 in reovirus platforms **(Fig S4F)**.

**Fig 6.**
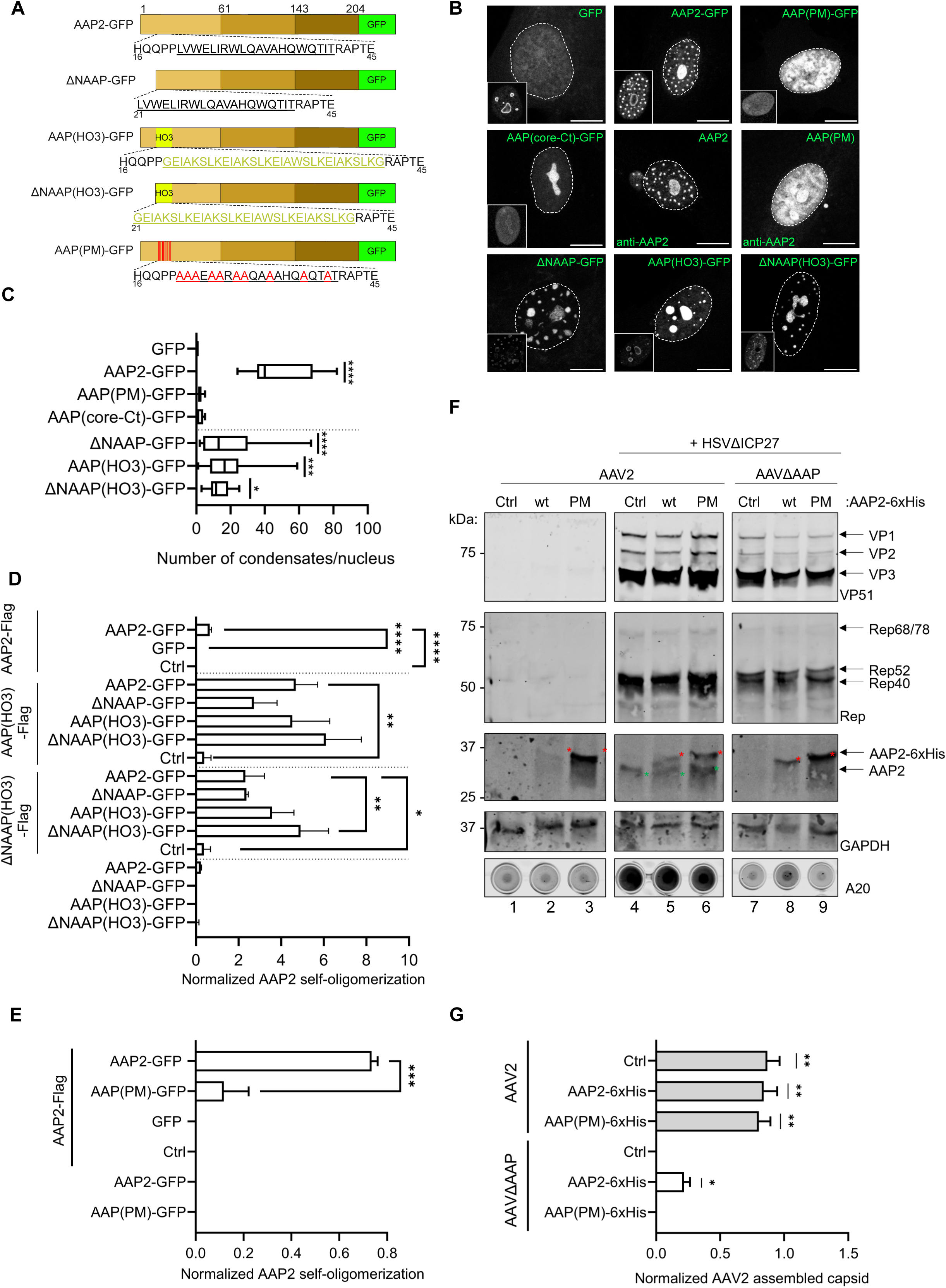
AAP2 hydrophobic residues in the amino acid region 21-40 are required for liquid-liquid phase separation and to trigger capsid assembly. (A) Schematic representation of the diverse AAP2 fused to GFP having i) deleted the N-terminal region (amino acids 1 to 20), ΔNAAP-GFP; ii) replaced predicted α-helix (amino acids 21 to 40) by HOTag3, AAP(HO3)-GFP; iii) deleted the N-terminal region and replaced predicted α-helix by HOTag3, ΔNAAP(HO3)-GFP and iv) substituted hydrophobic residues in predicted α-helix by alanines, AAP(PM)-GFP. The sequence of HOTag3 is indicated in yellow. The amino acid substitutions to alanines in AAP(PM)-GFP are indicated in red. The amino acids present in the predicted α-helix are underlined. (B) Immunofluorescence images of BJ cells expressing the AAP2-GFP, AAP2, and indicated mutants. At 16 hpt, the cells were fixed and immunostained for detection of nucleophosmin (anti-nucleophosmin, inset). The nuclei were stained with DAPI (dashed white line). For the detection of AAP2 and AAP(PM), the cells were immunostained with anti-AAP (white). The frame in the left bottom corner corresponds to NPM staining. The scale bar is 10 μm. (C) Plot of the numbers of nuclear AAP2-GFP condensates. The data represent the mean± SD; one-way ANOVA, (****)p< 0.0001, n ≥ 15. (D) Plot for the detection of the association of AAP2-GFP full-length or its indicated deletion mutants to AAP2-Flag, AAP(HO3)-Flag, or ΔNAAP(HO3)-Flag. The positive association was detected using a monoclonal anti-Flag antibody followed by incubation with anti-mouse conjugated to HRP. The data represents the mean ± SD of three independent experiments; one-way ANOVA, (*)p<0.01, (**)p<0.01, (***), p<0.001. (E) Plot for the detection of the association of AAP2-GFP full length or AAP(PM)-GFP to AAP2-Flag. The positive association was detected using a monoclonal anti-Flag antibody followed by incubation with anti-mouse conjugated to HRP. The data represents the mean ± SD of three independent experiments; one-way ANOVA, (***), p<0.001. (F) Immunoblotting (top) and dot blot of AAV2 progeny (bottom) of Vero cells co-infected with HSVΔICP27 (MOI 0.75) and either AAV2 (lanes 1-6) or AAVΔAAP (lanes 7 to 9)(MOI 10000). At 1 hpi, cells were transfected with pCI-Neo (Ctrl, lanes 1, 4, and 7), pCI-AAP2-6xHis (wt, lanes 2, 5, and 8), or pCI-AAP(PM)-6xHis (PM, lanes 3, 6, and 9). For immunoblotting, the cell extracts were harvested at 24 hpi, and the membrane was incubated for detection of the non-assembled capsid (anti-VP51), Rep proteins (anti-Rep), AAP2/AAP-6xHis (anti-AAP) and loading control (anti-GAPDH). For the dot blot of the virus progeny, the cells were harvested at 96 hpi, and the membrane was stained for detection of the assembled capsid (anti-A20 antibody). Red and green stars indicate the position of AAP2-6xHis and AAP2 variants, respectively. (G) Plot of normalized AAV2 assembled capsids rescued with wild-type or PM AAP2-6xHis of Vero cells co-infected with HSVΔICP27 and either AAV2 (grey bars) or AAVΔAAP (white bars). The data represent the mean ± SEM of three independent experiments; one-sample t-test with a hypothetical mean value of 0, (*) p< 0.05 and (**) p< 0.01.

We selected AAP(PM)-GFP for the following experiments since it has the most robust opposite phenotype to AAP2-GFP, and we built additional AAP2-GFP mutants to validate the role of the predicted AAP2 α-helix as follows (**Fig 6A**): *i)* ΔNAAP-GFP, with AAP2 residues 1 to 20 deleted, *ii)* AAP(HO3)-GFP, with the AAP2 predicted α-helix replaced by the well-characterized homo-hexameric HOTag3 (HO3) [47], and *iii)* ΔNAAP(HO3)-GFP, with residues 1 to 20 deleted and the predicted α-helix replaced by HOTag3. Notably, HOTag3 provides multivalency to proteins through its ability to form homo-hexamer structures [47]. After confirming their expression in WB (**Fig S3B, C, and E**), the AAP2-GFP mutants were compared to AAP2-GFP concerning their ability to form BCs. Interestingly, we found that similar to wild-type AAP2-GFP, the mutants ΔNAAP-GFP, AAP(HO3)-GFP, and ΔNAAP(HO3)-GFP successfully induced BC formation in BJ cells (**Fig 6B and C**).

Next, we investigated the self-oligomerization properties of the different AAP2-GFP mutants using the above-described AAP2-Flag/AAP2-GFP oligomerization assay **(Fig 5C**). In these conditions, both AAP(HO3)-Flag and ΔNAAP(HO3)-Flag were able to self-oligomerize with AAP2-GFP, ΔNAAP-GFP, AAP(HO3)-GFP, or ΔNAAP(HO3)-GFP **(Fig 6D)**. Interestingly, the replacement of the AAP2 α-helix by a HOTag3 significantly increased AAP2 self-oligomerization regardless of its binding partner. The deletion of the N-terminal region (aa 1-20) did not affect AAP2 self-oligomerization. In contrast, AAP(PM)-GFP had a significant decrease in its association with AAP2-Flag compared to wild-type AAP2-GFP (**Fig 6E**).

We then interrogated whether AAP2 self-oligomerization is necessary for AAV2 capsid assembly. To address this question, Vero cells expressing AAP2-6xHis or AAP(PM)-6xHis were infected with either AAV2 or recombinant AAVΔAAP, which contains a start-loss mutation for AAP2 but is silent for cap proteins. Simultaneously, the cells were co-infected with HSVΔICP27, an ICP27-deleted, replication-defective recombinant HSV-1. Thisvirus does not induce extensive cytopathic effect but still supports AAV2 replication [48, 49]. The co-infection of AAV2 with the helpervirus was confirmed by immunoblotting for the detection of disassembled AAV2 capsid (anti-VP51 antibody) and Rep proteins (**Fig 6F, lanes 4 to 9**). These AAV proteins are expressed only in co-infection with a helper virus. Similarly, the expression of AAP2-6xHis, AAP2(PM)-6xHis, and viral AAP2 were detected with anti-AAP antibody. As expected, AAP2 is not detected in the absence of a helper virus (**Fig 6F, lanes 1 to 3**) or upon co-infection of AAVΔAAP with HSVΔICP27 (**Fig 6F, lanes 7 to 9**). The assembly of wild-type AAV2 capsids was indeed unaffected by the presence of AAP2-6xHis or AAP(PM)-6xHis (**Fig 6F, lanes 4 to 6, and Fig 6G**), as assessed by dot blot of cell lysates incubated with an antibody recognizing assembled capsids (anti-capsid clone A20 antibody). However, in co-infection with HSVΔICP27, AAVΔAAP in presence of AAP2-6xHis showed capsid assembly, whereas AAP(PM)-6xHis did not, resulting in no rescue of assembled progeny virus. This outcome emphasizes the importance of AAP2 self-oligomerization for LLPS and AAV2 capsid assembly.

## Discussion

AAV biology has been extensively studied since its discovery as a contaminant of an AdV preparation [50]. Its recent importance stems from its potential as a gene therapy vector [1]. A deeper understanding of the AAV life cycle is essential to overcome the current obstacles, such as high enough production yield for the large doses needed [51], where an essential aspect relates to the functional capsid assembly mechanism.

The icosahedral AAV2 capsid assembly relies on the formation of three distinct capsid subunit interactions [52, 53]. While the interactions at the 2-fold axis are relatively straightforward, those at the 3- and 5-fold axes are more constrained and require additional assistance during capsid assembly [16]. While AAP2 has been shown to assist AAV capsid assembly [14], the mechanism responsible is unknown.

A large number of cellular BCs have been identified [37, 43, 54–56]. Similarly, many viruses can induce the formation of specialized intracellular compartments with LLPS properties [20–26]. In this study, we demonstrated that AAP2 fused to autofluorescent proteins possesses all the hallmarks of LLPS, which include i) large disordered regions, ii) globular morphology, iii) miscibility of the AAP2 inclusions, iv) dissolution after treatment to aliphatic diol treatment, v) recovering from photobleaching and vi) AAP2 inclusion size to be concentration-dependent. Therefore, AAP2 inclusion can be considered as proper BCs. We generated an AAP-specific antibody that, for the first time, revealed the BC properties of wild-type AAP2 in cells co-infected with AAV2 and a helper virus. This suggests a crucial role for AAP2 BCs in the AAV2 life cycle. The AAP-specific antibody was generated in rabbits that were immunized with the peptide SARRIKDASRRSQQTS from AAP2, which is partially conserved in AAV serotypes 1, 3B, 6, 7, and 9. Therefore, this antibody may be used to visualize AAP BCs in these specific AAV serotypes as well.

Interestingly, AAP2 forms two kinds of BCs, nucleolar and small extranucleolar. The small extranucleolar AAP2 condensates co-localize with NPM but not with fibrillarin, which are both nucleolar markers. NPM can form BCs by itself or localize to nucleoli via multivalent interactions with proteins containing arginine-rich linear motifs [57]. AAP2 contains 10% arginine in its primary sequence with an arginine linear motif between residues 147 to 157, suggesting a potential site for association with NPM. These results are also consistent with previous observations describing significant co-localization of NPM and AAV2 cap proteins [58]. These data also align with the earlier observationsfrom Earley et al., 2017 [15], which described the localization of nucleostemin in both types of AAP2 BCs. Additionally, both NPM and nucleostemin have been shown to interact directly within the nucleolus [59] and to shuttle proteins between the nucleolus and the cytoplasm [60, 61]. This suggests that AAP2 BCs may act as protein hubs that facilitate assembly coordination.

Still, the role of AAP LLPS in the AAV life cycle remains unclear. LLPS may enable the concentration of capsid proteins in a localized area at sufficient density to trigger the capsid assembly mechanism. However, simply concentrating VP3 in the nucleolus by adding a nucleolar localization signal is insufficient to trigger capsid assembly [14], suggesting the involvement of an additional mechanism. Some virus proteins induce the formation of LLPS to provide a scaffolding function that aids the assembly reaction, as is hypothesized for the HSV-1 UL11 protein, which is thought to provide a bridging function for UL16 [62, 63]. Similarly, the AAP LLPS properties may provide a flexible scaffold providing the proper conditions for assembling the AAV capsid proteins. Furthermore, viral LLPS compartments can also function as a shield, preventing unwanted host factors from entering. In this context, we observed that fibrillarin did not colocalize with the extranucleolar compartments formed upon AAP2 expression alone, unlike NPM and nucleostemin. Fibrillarin is a protein from the dense fibrillar component in the nucleolus, capable of methylating both RNA [64] and proteins [65]. Fibrillarin function might be unwanted in AAP2 BC, potentially inhibiting AAV2 capsid assembly. This is supported by data showing that AAV2 capsid uncoating occurs in the presence of fibrillarin [48].

We also demonstrated through screening of AAP2 deletion mutants that the N-terminal region (amino acids 1 to 60) contains all the elements for AAP2 BC formation, including the ability to self-oligomerize. According to PONDR prediction, the AAP2 N-terminal region is partially ordered, which is consistent with the AlphaFold2 prediction for the presence of an *α*-helix spanning AAP2 residues 21 to 40. In fact, this region contains hydrophobic residues exposed at one side of the *α*-helix (**Fig S4A i**). In general, the *α*-helix is a not-so-stable secondary structure but becomes more stable when coiled-coil helices are generated through association with other *α*-helices [66]. Bulky hydrophobic amino acids play an important role in protein-protein interactions and oligomerization due to their ability to form stable hydrophobic interactions [67]. AlphaFold2 was able to predict AAP2 oligomerization **(Fig S4A ii)**. Indeed, the AAP2 region from amino acids 21-40 contains three leucine residues, which are known to promote the formation of *α*-helices through non-covalent H-bond interactions [68]. Also, this region contains three tryptophans, which have been described to stabilize the coiled-coil oligomerization of an *α*-helix when arranged in the correct disposition [69]. AAP(PM) was specifically designed to not interfere with the *α*-helix structure in AlphaFold2 prediction **(Fig S4A iii)** but to reduce aliphatic interaction through hydrogen bonds. On this basis, AlphaFold2 could not predict AAP(PM) self-oligomerization (data not shown). We experimentally assessed the presence of this predicted AAP2 *α*-helix and demonstrated with AAP(PM)-GFP that the bulky hydrophobic residues are required to form AAP2 BCs and support self-oligomerization. Moreover, the replacement of the predicted AAP2 *α*-helix by an HOTag3 [70], which provides multivalency through homo-hexamers, allowed AAP(HO3) formation of BCs and self-oligomerization. Notably, the deletion of amino acids 1 to 20 in ΔNAAP-GFP and ΔNAAP(HO3)-GFP did not disturb their ability to self-oligomerize and form extranuclear condensates. Collectively, these data support the hypothesis that AAP2 possesses an *α*-helix between residues 1 to 40 which is required for self-oligomerization, BC formation, and capsid assembly. The mutation of the bulky hydrophobic amino acids disrupted self-oligomerization and prevented BC formation and AAV2 capsid assembly.

Among the 13 AAV serotypes, AAP has a high conservation ratio [71], particularly in its N-terminal region, including the AAP α-helix. A specific example is AAP1 [18], where replacing the core and C-terminal regions with a generic linker containing nucleolar localization signals still retains its ability to support capsid assembly, suggesting that this role is also present in the AAP1 N-terminal region.

In summary, this study shows that AAP2 BC formation and self-oligomerization rely on bulky residues in the α-helix at its N-terminus. These residues are essential for AAV capsid assembly, as their mutation significantly inhibits progeny virus formation. Identifying the crucial amino acids of AAP involved in assembly and LLPS could aid in targeted AAV capsid mutagenesis, potentially enhancing efficiency and tissue targeting of AAV vectors in gene therapy.

## Material and Methods

### Cells and Viruses

BJ cells (human foreskin BJ fibroblasts, ATCC CRL-2522™) and Vero cells (African green monkey kidney, ATCC CCL-81) were obtained from ATCC (Manassas, Virginia, USA) and cultured in Dulbecco’s Modified Eagle Medium (DMEM) supplemented with 10% fetal calf serum (FCS), 100 μg/ml streptomycin and 100 units/ml penicillin (P/S) in a humified incubator at 37°C and 5% CO_2_.

Purified wtAAV2 was produced by the Viral Vector Facility (VVF) of the Neuroscience Center Zurich (ZNZ) (University of Zurich, Switzerland) as previously described [72, 73] using the pAV2 plasmid [74]. The recombinant vector of AAV serotype 2 (AAVΔAAP), containing a mutation in the AAP start codon (CTG→CGG) in pAV2(ΔAAP) was produced by the VVF of the ZNZ. AdV-5 was kindly provided by Dr. Urs Greber and Dr. Marit Suomalainen (Dept of Molecular Life Sciences University of Zurich, Switzerland). The stocks of wild-type HSV-1 (strain F, kindly provided by Dr. B. Roizman, University of Chicago, USA) and HSVΔICP27 (kindly provided by Dr. R. Everett, University of Glasgow, UK) were amplified and purified as previously described [75].

### Antibodies

The following antibodies were purchased: mouse monoclonal anti-AAV2 intact particle (clone A20, ProGen), rabbit anti-AAV VP1/VP2/VP3 (VP51, ProGen), mouse anti-AAV2 Rep (10R-A111A, Fitzgerald Industries), mouse anti-α-tubulin (B-5-1-12, Merck), mouse anti-β-actin (AC-74, Merck), rabbit anti-vimentin (H-84, Santa Cruz), mouse anti-GAPDH antibody (6C5, Abcam), mouse anti-GFP (B-2, Santa Cruz), rabbit anti-fibrillarin (ab5821, Abcam), mouse anti-nucleophosmin (FC82291, Abcam), mouse anti-Flag (M2, Merck).

The rabbit polyclonal AAP antiserum (C-SARRIKDASRRSQQTS) was produced and affinity purified by Pacific Immunology® (Ramona, CA, USA).

### DNA plasmids

AAP2 was PCR amplified from pAV2 (ATCC 37216)[76] using specific primers to insert *Xho*I and *Mlu*I sites at the 5’ and 3’ ends of AAP2, followed by ligation on those sites in pCI-Neo (Promega) to obtain pCI-AAP2 non-stop. pCI-AAP2-GFP was obtained by PCR amplification of pCI-EGFP [46] using specific primers to insert *Mlu*I site in-frame at 5’end and stop codon/*Not*I site at 3’end of EGFP, followed by ligation on those sites in pCI-AAP2 non-stop. pCI-AAP2-mKO2 was obtained by PCR amplification of pFug-Fucci-G0/1(MBL Life science, Japan) using specific primers to insert in-frame an *Xba*I site at 5’ end and stop codon/*Not*I site at 3’end of mKO2, followed by ligation on those sites in pCI-AAP2 non-stop. pCI-AAP2-mCherry-(471-721)µNS was obtained by PCR amplification of AAP2 of pCI-AAP2-GFP using specific primers to insert *Xho*I and *MluI*, followed by ligation in-frame on those in pCI-mCherry-(471-721)µNS [46]. AAP Nt, core, and Ct deletion mutant primers were produced by amplifying desired sequences using specific primers to add *Nhe*I and *Mlu*I sites, followed by in-frame ligation in those sites in pCI-AAP2-GFP.

For the production of pCI-ΔNAAP-GFP, pCI-AAP(HO3)-GFP, pCI-ΔNAAP(HO3)-GFP, and pCI-AAP(PM)-GFP, synthetic gene fragments harboring at 5’ and3’ ends *Nhe*I and *Ava*I sites were produced at Eurofins (**Table S2**) cloned into pCI-AAP2-GFP. pCI-AAP2-6xHis, pCI-AAP2-Flag, and respective mutants were produced by annealing oligonucleotides with 6xHis or Flag sequences with *Mlu*I and *Not*I overhangs followed by ligation on those sites into pCI-AAP2-GFP. pCI-AAP2-stop and pCI-AAP(PM)-stop were produced by annealing primers with a stop codon with *Mlu*I and *Not*I overhangs and ligated on those sites in pCI-AAP2-GFP and pCI-AAP(PM)-GFP plasmids, respectively.

The oligonucleotides were synthesized at Microsynth (Switzerland) and described in **Table S1**.

### IDR prediction

The intrinsically disordered regions of AAP2 were determined using PONDR® (Molecular Kinetics, Inc, http://www.pondr.com) using the VSL2 algorithm. Data were plotted with GraphPad Prism (version 10.1.1(270)).

### Transfection

24 h prior to transfection, BJ cells were seeded at a density of 3×10^4^ cells per well onto 24-well tissue culture plates. The cells were transfected with 750 ng DNA, or otherwise indicated in the figure legends, mixed 1 μl Plus™ reagent (Thermo Fisher Scientific) in 40 μl of OptiMEM (Thermo Fisher Scientific), and incubated for 8 min at room temperature. Meanwhile, 1.7 μl of Lipofectamine™ LTX transfection reagent (Thermo Fisher Scientific) was mixed with 25 μl of OptiMEM and added to the DNA-Plus Reagent mixture followed by 15 min incubation at room temperature. The Cells were washed twice with OptiMEM, supplemented with 150 μl OptiMEM, and the transfection mix was added dropwise on top.

Similarly, Vero cells were plated at a density of 1×10^5^ cells per well onto 24-well plates 24 h prior to transfection. For transfection, 750 ng of DNA (or otherwise specified amount) was mixed in 50 μl of OptiMem. Separately, 3 μl of Lipofectamine™ 2000 transfection reagent (Thermo Fisher Scientific) was mixed with 50 μl of OptiMEM. The Lipofectamine mixture was then added to the DNA mix and incubated for 20 min at room temperature. Following the incubation, the cells were washed twice with PBS (phosphate buffered saline: 1 mM KH2PO4, 3 mM Na_2_HPO_4_-7H_2_0, 155 mM NaCl at pH 7.4), supplemented with 400 μl of 10% FCS in DMEM and the transfection mixture was added dropwise onto the cells.

### Immunofluorescence

Immunofluorescence was conducted as described previously [36]. Briefly, the cells were fixed with 2% paraformaldehyde in PBS for 10 min at room temperature, followed by three washes with PBS. Subsequently, the cells were permeabilized with 0.1% Triton-X 100 in PBS for 10 min, washed thrice with PBS, and blocked with 1% BSA in PBS for 20 min. Then, the cells were incubated for 45 min in a humid chamber with the primary antibody, washed thrice with PBS, and then incubated for 45 min with the secondary antibody along with 1 μg/ml of DAPI (4′,6-diamidino-2-phenylindole). The samples were mounted using ProLong™ Gold antifade mountant (ThermoFisher Scientific). The images were acquired using a Leica SP8 (DM550Q) confocal laser scanning microscope (CLSM). The data and statistical analyses were performed using Fiji software [77] (version: 2.13.0/1.54f; http://imageJ.net/Contributors) and GraphPad Prism, version 10.1.2 for Windows (GraphPad Software, Boston, MA, USA, www.graphpad.com), respectively.

### Immunoblotting

In general, the cells were lysed by adding 25 µl of 4x Laemmli sample buffer (8% SDS, 40% glycerol, 200 mM Tris-HCl pH 6.8, and 0.4% bromophenol blue and 142 mM 2-mercaptoethanol). The cell extracts were heated for 10 min at 95°C, sonicated for 5 seconds at 14 Hz, and loaded onto an SDS-polyacrylamide gel. The proteins were separated at 30 mA per gel and transferred to a 0.45 µm Protran nitrocellulose membrane (Amersham). The membranes were blocked for 30 min in 5% milk-PBS, followed by incubation with primary and the corresponding secondary antibodies conjugated to either IRdye® 680 or 800 (LI-COR). The membrane was acquired using the Odyssey® M Imaging System and Empiria Studio® Software (LI-COR Biosciences).

### Quantification of number, area, and solidity of AAP2 condensates

The AAP condensates number, area, and solidity were obtained using the “analyze particles tool” from Fiji software [77] (version: 2.13.0/1.54f; http://imageJ.net/Contributors), as previously described [36, 78]. The data were processed using Microsoft Excel® for Microsoft 365 MSO (Version 2406). The Statistical analysis, unpaired parametric two-ANOVA, and plots were performed using Prism 10 for Windows 64-bit (version 10.2.3 (403)) (GraphPad Software, LLC).

### Determination of AAP2 phase diagram

BJ cells were seeded over coverslip at 3×10^4^ cells per well onto a 24-well tissue culture plate. At 24 h, the cells were transfected with Lipofectamine LTX transfection reagent as indicated above, with increasing amounts of DNA plasmid (250, 500, 750, 1000, 1250, and 1500 ng/well). At 16 hpt, cells were fixed and prepared for immunoblotting and immunofluorescence as described above. After the quantification of BCs, the datasets were graphed and used to compute second-order polynomial regression curves, followed by determining the first derivative of the phase diagram to identify the DNA concentration that leads to maximum phase separation. The calculations were performed employing Prism 10 for Windows 64-bit (version 10.2.3 (403)) (GraphPad Software, LLC).

### 1,6-Hexanediol treatment

At the indicated time, the cell medium was replaced by a medium containing 3.5% 1,6-hexanediol (Merck) in 2% FCS-DMEM. Subsequently, cells were incubated at 37°C for 5 min and then fixed with 2% paraformaldehyde for 10 min at room temperature. The cells were immunostained as described above.

### FRAP

FRAP measurements were performed as described previously [36]. Briefly, 2×10^3^ BJ cells per well were seeded in μ-Slide 18-well glass-bottom plates (Ibidi). The cells were transfected with Lipofectamine LTX and pCI-AAP-mKO2, as described above. At 16 hpt, the nuclei were stained with 1 μg/ml Hoechst 33342 diluted in FluoroBRITE DMEM (Gibco), incubated for 30 min at 37°C/5% CO_2_, and subjected to FRAP analysis. FRAP measurements were performed with a Leica SP8 Falcon CLSM using the FRAP function of the LasX software (Leica) at 37°C, 5% CO_2_, 95% humidity as follows: a circular area of 1.5 μm in diameter encompassing condensates either >, =, or < than 1.5 μm in diameter was bleached at 100% laser power, for five iterations. The fluorescent recovery was monitored by taking fluorescence images for 45 iterations. As the fluorescent control for each FRAP acquisition, a circular area of 1.5 μm was used, and a squared area (3 μm x 3 μm) outside of a cell was chosen as the background. The FRAP data set was analyzed using easyFRAP-web [79]. The fully normalized data were used to generate FRAP diagrams and calculate the recovery half-times (T-half) and mobile fractions from independent measurements.

### Live cell imaging

For this purpose, 2×10^3^ BJ cells were plated in μ-Slide 18-well glass-bottom plates. The cells were then transfected with pCI-AAP-mKO2, as described above. At 16 hpt, the nuclei were stained using 1 μg/mL Hoechst 33342 diluted in FluoroBRITE DMEM (Gibco) for 30 min at 37°C. The samples were imaged using a Leica SP8 Falcon CLSM set at 37°C, 5% CO_2_, and 95% humidity.

### Co-infection of AAV with helper viruses

One day prior to infection, Vero cells were seeded at a density of 1×10^5^ cells per well in a 12-well tissue culture plate. The medium was exchanged with virus inoculum (specific viruses and their corresponding MOI are described in the figure legends) to initiate infection. Following a 30-minute adsorption period at 4°C while shaking, the cells were transferred to a humidified incubator set at 37°C and 5% CO_2_ for 1 hour. Subsequently, the virus inoculum was replaced with DMEM supplemented with 2% FCS, and the cells were further incubated for the indicated time points.

### Dot blot assay for detection of AAV progeny

The day before infection, Vero cells were seeded at 1×10^5^ cells per well in 12-well tissue culture plates. The next day, cells were infected, as described above. Briefly, virus dilution was added in 100 μl DMEM-SF, and plates were incubated at 4°C shaking for 30 min. Then, the plates were transferred to 37°C. Meanwhile, the transfection mixture was prepared in OptiMEM using Lipofectamine™ 2000. After 1 h incubation, 600 μl DMEM-2%FCS without antibiotics was added, and the transfection mixture was added dropwise on top. At 8 hpi, the inoculum was removed, the cells were washed carefully with PBS, and the medium was replaced with 500 μl DMEM-2%FCS without antibiotics. At 24 hpi, the samples from one plate were collected for immunoblotting analysis. At 96 hpi, the cells and supernatants of the second plate were collected. For this, the plate was freeze-thawed for three cycles of liquid nitrogen and 37°C, followed by spinned down at 15’000 x g for 7 min at 4°C. Then, opaque filter plates (MultiSreen HTS IP filter plate, 0.45 μm hydrophobic PVDF, Millipore-Merck, Cat # MSIPS4W10) were activated for 1 min with methanol and washed twice with PBS, using the MultiScreen HTS Vacuum Manifold (Millipore, MSVMHTS00). The virus supernatant was loaded and filtered. The membrane was washed twice with PBS, blocked twice for 30 min with 5% BSA in PBS, and then incubated overnight at 4°C with the indicated primary antibody. The plate was washed three times for 2 min with 5% BSA in PBS and incubated for 1 h at room temperature with corresponding secondary antibody conjugated to IRDye^®^ 800 (LI-COR). The plate was washed three times with PBS. The data were acquired with Odyssey^®^ M Imaging System and analyzed using Empiria Studio^®^ software. Statistical analysis was performed using GraphPad Prism, version 10.1.2 for Windows (GraphPad Software, Boston, MA, USA, www.graphpad.com).

### In vivo protein-protein interaction assay

The in vivo protein-protein assay was performed as described previously [45, 46, 80]. Briefly, BJ cells were seeded and 24 h later transfected with Lipofectamine LTX transfection reagent, as indicated above, with 750 ng of «fish» protein and 750 ng of «bait» protein. At 16 hpt, cells were fixed and prepared for immunofluorescence as outlined above.

### Capture ELISA for AAP2 self-oligomerization

The day before transfection, Vero cells were seeded at a density of 1.3×10^5^ cells per well in 12-well tissue culture plates. The cells were transfected according to the manufacturer’s instructions. At 24 hpt, the cells were harvested using RIPA lysis buffer (50 mM Tris HCl, pH 7.4, 150 mM NaCl, 1 mM EDTA, 1 mM EGTA, 0.5% Triton X-100, and 0.5% sodium deoxycholate), supplemented with EDTA-free protease inhibitor cocktail (Roche). The cell lysates were clarified by centrifugation at 15’000 x g for 7 min at 4°C. ELISA MaxiSorp immunoplates (Thermo Fisher Scientific) were prepared by washing once with 200 µl coating buffer (50 mM carbonate/bicarbonate buffer pH 9.6) and then coated with 0.5 µg/ml polyclonal anti-GFP antibody (ab6556, Abcam) diluted in 100 µl coating buffer for 24 h at 4°C. At room temperature, the plate was blocked with 1% BSA in PBS-T for 1 h. Next, 100 µl cell lysate diluted in PBS was added per well and incubated for 16 h at 4°C. The plate was then incubated for 1 h at room temperature with 50 µl mouse anti-Flag(clone M2, Sigma) diluted 1: 1000 in 5% milk-PBS. The plates were incubated for 1 h at room temperature with 100 µl anti-mouse conjugated to HRP (Sigma) diluted 1:5000 in 5% milk-PBS. The plates were washed between each incubation step three times with 200 µl PBS-T (PBS containing 0.05% Tween 20) followed by 2 min shaking in the orbital shaker. The reaction was developed using 50 µl of Pierce TMB substrate kit (ThermoFisher) and incubated for 30 min at room temperature in the dark. The reaction was stopped with 50 µl of 1M H_2_SO_4_. The absorbance was measured at 450 nm using a Tecan plate reader Infinite F50. The normalized Flag value was calculated using this formula: Normalized Flag value = (“Flag signal” – “AAP2-Flag”)/“AAP2-GFP/AA2P-Flag.” In this formula, “Flag signal” is the absorbance value for the condition to be normalized, “AAP2-Flag” is the absorbance value of the AAP2-Flag condition, and “AAP2-GFP/AAP2-Flag” is the absorbance value of the co-expression of AAP2-GFP and AAP2-Flag. Statistical analysis was performed using GraphPad Prism version 10.2.0 for Windows, GraphPad Software, Boston, Massachusetts USA, www.graphpad.com.

### AlphaFold2 predictions

Protein structures of AAP2 N-terminus monomer, AAP2 pentamer, and AAP(PM)-N-terminus monomer were predicted using AlphaFold2 Server (https://colab.research.google.com/github/sokrypton/ColabFold/blob/main/AlphaFold2.ipynb) [81].

## Supporting information

suppl Fig S1, S2,S3, S4 and corresponding Fig Legends and Tables S1 and S2

## Author contributions

The authors declare no conflict of interest.

Conceptualization: J.V., C.E. and C.F.; Methodology: J.V., M.K. and C.E.; Software: J.V. and C.E.; Validation: J.V. and C.E.; Formal analysis: J.V. and C.E.; Investigation: J.V. and C.E.; Resources: C.E. and C.F.; Data Curation: J.V. and C.E.; Writing-Original Draft: J.V. and C.E.; Writing-Review & Editing: J.V., C.E. and C.F.; Visualization: J.V. and C.E.; Supervision: C.E. and C.F.; Project Administration: C.E. and C.F.; Funding Acquisition: C.F.

## References

1. Wang D, Tai PWL, Gao G. Adeno-associated virus vector as a platform for gene therapy delivery. Nature Reviews Drug Discovery 2019 18:5. 2019;18(5):358–78.

2. Meier AF, Fraefel C, Seyffert M. The interplay between adeno-associated virus and its helper viruses. Viruses: MDPI AG; 2020.

3. Samulski RJ, Zhu X, Xiao X, Brook JD, Housman DE, Epstein N, et al. Targeted integration of adeno-associated virus (AAV) into human chromosome 19. The EMBO Journal. 1991;10(12):3941–50-50.

4. Kotin RM, Siniscalco M, Samulski RJ, Zhu XD, Hunter L, Laughlin CA, et al. Site-specific integration by adeno-associated virus. Proceedings of the National Academy of Sciences. 1990;87(6):2211–5.

5. Schnepp Bruce C, Jensen Ryan L, Chen C-L, Johnson Philip R, Clark KR. Characterization of Adeno-Associated Virus Genomes Isolated from Human Tissues. Journal of Virology. 2005;79(23):14793–803.

6. Wistuba A, Kern A, Weger S, Grimm D, Kleinschmidt JA. Subcellular compartmentalization of adeno-associated virus type 2 assembly. Journal of virology. 1997;71(2):1341–52.

7. Srivastava A, Lusby EW, Berns KI. Nucleotide sequence and organization of the adeno-associated virus 2 genome. Journal of Virology. 1983;45(2):555–64.

8. Lusby E, Fife KH, Berns KI. Nucleotide sequence of the inverted terminal repetition in adeno-associated virus DNA. Journal of Virology. 1980;34(2):402–9.

9. Laughlin CA, Westphal H, Carter BJ. Spliced adenovirus-associated virus RNA. Proceedings of the National Academy of Sciences. 1979;76(11):5567–71.

10. Trempe JP, Carter BJ. Alternate mRNA splicing is required for synthesis of adeno-associated virus VP1 capsid protein. Journal of Virology. 1988;62(9):3356–63.

11. Samulski RJ, Muzyczka N. AAV-Mediated Gene Therapy for Research and Therapeutic Purposes. Annual Review of Virology. 2014;1(1):427–51.

12. Galibert L, Hyvönen A, Eriksson RAE, Mattola S, Aho V, Salminen S, et al. Functional roles of the membrane-associated AAV protein MAAP. Scientific Reports. 2021;11:21698-.

13. Ogden PJ, Kelsic ED, Sinai S, Church GM. Comprehensive AAV capsid fitness landscape reveals a viral gene and enables machine-guided design. Science. 2019;366(6469):1139–43.

14. Sonntag F, Schmidt K, Kleinschmidt JA. A viral assembly factor promotes AAV2 capsid formation in the nucleolus. Proceedings of the National Academy of Sciences of the United States of America. 2010;107(22):10220–5.

15. Earley LF, Powers JM, Adachi K, Baumgart JT, Meyer NL, Xie Q, et al. Adeno-associated Virus (AAV) Assembly-Activating Protein Is Not an Essential Requirement for Capsid Assembly of AAV Serotypes 4, 5, and 11. Journal of virology. 2017;91(3):1–21.

16. Naumer M, Sonntag F, Schmidt K, Nieto K, Panke C, Davey NE, et al. Properties of the Adeno-Associated Virus Assembly-Activating Protein. Journal of Virology. 2012;86(23):13038–48.

17. Sonntag F, Kother K, Schmidt K, Weghofer M, Raupp C, Nieto K, et al. The Assembly-Activating Protein Promotes Capsid Assembly of Different Adeno-Associated Virus Serotypes. Journal of Virology. 2011;85(23):12686–97.

18. Tse LV, Moller-Tank S, Meganck RM, Asokan A. Mapping and Engineering Functional Domains of the Assembly-Activating Protein of Adeno-associated Viruses. Journal of Virology. 2018;92(14):1–18.

19. Earley LF, Kawano Y, Adachi K, Sun X-X, Dai M-S, Nakai H. Identification and Characterization of Nuclear and Nucleolar Localization Signals in the Adeno-Associated Virus Serotype 2 Assembly-Activating Protein. Journal of Virology. 2015;89(6):3038–48.

20. Zhou Y, Su Justin M, Samuel Charles E, Ma D. Measles Virus Forms Inclusion Bodies with Properties of Liquid Organelles. Journal of Virology. 2019;93(21):10.1128/jvi.00948-19.

21. Heinrich BS, Maliga Z, Stein DA, Hyman AA, Whelan SPJ. Phase transitions drive the formation of vesicular stomatitis virus replication compartments. mBio. 2018;9(5):1–10.

22. Nikolic J, Le Bars R, Lama Z, Scrima N, Lagaudrière-Gesbert C, Gaudin Y, et al. Negri bodies are viral factories with properties of liquid organelles. Nature Communications. 2017;8(1):1–12.

23. Caragliano E, Bonazza S, Frascaroli G, Tang J, Soh TK, Grünewald K, et al. Human cytomegalovirus forms phase-separated compartments at viral genomes to facilitate viral replication. Cell Reports. 2022;38(10):110469-.

24. Seyffert M, Georgi F, Tobler K, Bourqui L, Anfossi M, Michaelsen K, et al. The HSV-1 Transcription Factor ICP4 Confers Liquid-Like Properties to Viral Replication Compartments. International Journal of Molecular Sciences 2021, Vol 22, Page 4447. 2021;22(9):4447-.

25. Geiger F, Acker J, Papa G, Wang X, Arter WE, Saar KL, et al. Liquid–liquid phase separation underpins the formation of replication factories in rotaviruses. The EMBO Journal. 2021:e107711-e.

26. Chen H, Cui Y, Han X, Hu W, Sun M, Zhang Y, et al. Liquid–liquid phase separation by SARS-CoV-2 nucleocapsid protein and RNA. Cell Research: Springer Nature; 2020. p. 1-.

27. Li H, Ernst C, Kolonko-Adamska M, Greb-Markiewicz B, Man J, Parissi V, et al. Phase separation in viral infections. Trends in Microbiology. 2022.

28. Banani SF, Lee HO, Hyman AA, Rosen MK. Biomolecular condensates: Organizers of cellular biochemistry. Nature Reviews Molecular Cell Biology: Nature Publishing Group; 2017. p. 285–98.

29. Martin EW, Holehouse AS. Intrinsically disordered protein regions and phase separation: sequence determinants of assembly or lack thereof. Emerging Topics in Life Sciences. 2020;4(3):307–29.

30. Mohanty P, Kapoor U, Sundaravadivelu Devarajan D, Phan TM, Rizuan A, Mittal J. Principles Governing the Phase Separation of Multidomain Proteins. Biochemistry. 2022;61(22):2443–55.

31. Li J, Zhang M, Ma W, Yang B, Lu H, Zhou F, et al. Post-translational modifications in liquid-liquid phase separation: a comprehensive review. Molecular Biomedicine. 2022;3(1):13.

32. Van Treeck B, Parker R. Emerging Roles for Intermolecular RNA-RNA Interactions in RNP Assemblies. Cell. 2018;174(4):791–802.

33. Alberti S, Gladfelter A, Mittag T. Considerations and Challenges in Studying Liquid-Liquid Phase Separation and Biomolecular Condensates. Cell. 2019;176(3):419–34.

34. Brangwynne CP, Eckmann CR, Courson DS, Rybarska A, Hoege C, Gharakhani J, et al. Germline P Granules Are Liquid Droplets That Localize by Controlled Dissolution/Condensation. Science. 2009;324(5935):1729-32.

35. Fei J, Jadaliha M, Harmon TS, Li ITS, Hua B, Hao Q, et al. Quantitative analysis of multilayer organization of proteins and RNA in nuclear speckles at super resolution. Journal of Cell Science. 2017;130(24):4180–92.

36. Vetter J, Papa G, Seyffert M, Gunasekera K, Lorenzo GD, Wiesendanger M, et al. Rotavirus Spike Protein VP4 Mediates Viroplasm Assembly by Association to Actin Filaments. Journal of Virology. 2022;96(17):10.1128/JVI.01074-22.

37. Molliex A, Temirov J, Lee J, Coughlin M, Kanagaraj Anderson P, Kim Hong J, et al. Phase Separation by Low Complexity Domains Promotes Stress Granule Assembly and Drives Pathological Fibrillization. Cell. 2015;163(1):123–33.

38. Peskett TR, Rau F, O’Driscoll J, Patani R, Lowe AR, Saibil HR. A Liquid to Solid Phase Transition Underlying Pathological Huntingtin Exon1 Aggregation. Molecular Cell. 2018;70(4):588–601.e6.

39. Lee K-H, Zhang P, Kim HJ, Mitrea DM, Sarkar M, Freibaum BD, et al. C9orf72 Dipeptide Repeats Impair the Assembly, Dynamics, and Function of Membrane-Less Organelles. Cell. 2016;167(3):774–88.e17.

40. Elbaum-Garfinkle S. Matter over mind: Liquid phase separation and neurodegeneration. Journal of Biological Chemistry. 2019;294(18):7160–8.

41. Kroschwald S, Maharana S, Mateju D, Malinovska L, Nüske E, Poser I, et al. Promiscuous interactions and protein disaggregases determine the material state of stress-inducible RNP granules. eLife. 2015;4:e06807.

42. Axelrod D, Koppel DE, Schlessinger J, Elson E, Webb WW. Mobility measurement by analysis of fluorescence photobleaching recovery kinetics. Biophysical Journal. 1976;16(9):1055–69.

43. Shin Y, Brangwynne CP. Liquid phase condensation in cell physiology and disease. Science. 2017;357(6357).

44. Carter GC, Hsiung C-H, Simpson L, Yang H, Zhang X. N-terminal Domain of TDP43 Enhances Liquid-Liquid Phase Separation of Globular Proteins. Journal of Molecular Biology. 2021;433(10):166948.

45. Miller CL, Arnold MM, Broering TJ, Eichwald C, Kim J, Dinoso JB, et al. Virus-derived Platforms for Visualizing Protein Associations inside Cells*. 2007.

46. Nguyen CL, Eichwald C, Nibert ML, Münger K. Human Papillomavirus Type 16 E7 Oncoprotein Associates with the Centrosomal Component γ-Tubulin. Journal of Virology. 2007;81(24):13533–43.

47. Zhang Q, Huang H, Zhang L, Wu R, Chung C-I, Zhang S-Q, et al. Visualizing Dynamics of Cell Signaling In Vivo with a Phase Separation-Based Kinase Reporter. Molecular Cell. 2018;69(2):334–46.e4.

48. Sutter SO, Lkharrazi A, Schraner EM, Michaelsen K, Meier AF, Marx J, et al. Adeno-associated virus type 2 (AAV2) uncoating is a stepwise process and is linked to structural reorganization of the nucleolus. PLoS pathogens. 2022;18(7):e1010187-e.

49. Aubert M, Blaho John A. The Herpes Simplex Virus Type 1 Regulatory Protein ICP27 Is Required for the Prevention of Apoptosis in Infected Human Cells. Journal of Virology. 1999;73(4):2803–13.

50. Atchison RW, Casto BC, Hammon WM. Adenovirus-Associated Defective Virus Particles. Science. 1965;149(3685):754–6.

51. Colella P, Ronzitti G, Mingozzi F. Emerging Issues in AAV-Mediated *In Vivo* Gene Therapy. Molecular Therapy Methods & Clinical Development. 2018;8:87–104.

52. Agbandje-Mckenna M, Kleinschmidt J. Chapter 3 AAV Capsid Structure and Cell Interactions. Methods in Molecular Biology. 2011;807.

53. Xie Q, Bu W, Bhatia S, Hare J, Somasundaram T, Azzi A, et al. The atomic structure of adeno-associated virus (AAV-2), a vector for human gene therapy. Proceedings of the National Academy of Sciences. 2002;99(16):10405–10.

54. Feric M, Vaidya N, Harmon TS, Mitrea DM, Zhu L, Richardson TM, et al. Coexisting Liquid Phases Underlie Nucleolar Subcompartments. Cell. 2016;165(7):1686–97.

55. Larson AG, Elnatan D, Keenen MM, Trnka MJ, Johnston JB, Burlingame AL, et al. Liquid droplet formation by HP1α suggests a role for phase separation in heterochromatin. Nature. 2017;547(7662):236–40.

56. Brangwynne CP, Mitchison TJ, Hyman AA. Active liquid-like behavior of nucleoli determines their size and shape in Xenopus laevis oocytes. Proceedings of the National Academy of Sciences. 2011;108(11):4334–9.

57. Mitrea DM, Cika JA, Guy CS, Ban D, Banerjee PR, Stanley CB, et al. Nucleophosmin integrates within the nucleolus via multi-modal interactions with proteins displaying R-rich linear motifs and rRNA. Elife. 2016;5.

58. Bevington JM, Needham PG, Verrill KC, Collaco RF, Basrur V, Trempe JP. Adeno-associated virus interactions with B23/Nucleophosmin: Identification of sub-nucleolar virion regions. Virology. 2007;357(1):102–13.

59. Ma H, Pederson T. Nucleophosmin is a binding partner of nucleostemin in human osteosarcoma cells. Mol Biol Cell. 2008;19(7):2870–5.

60. Tsai RYL, McKay RDG. A multistep, GTP-driven mechanism controlling the dynamic cycling of nucleostemin. Journal of Cell Biology. 2005;168(2):179–84.

61. Borer RA, Lehner CF, Eppenberger HM, Nigg EA. Major nucleolar proteins shuttle between nucleus and cytoplasm. Cell. 1989;56(3):379–90.

62. Metrick Claire M, Koenigsberg Andrea L, Heldwein Ekaterina E. Conserved Outer Tegument Component UL11 from Herpes Simplex Virus 1 Is an Intrinsically Disordered, RNA-Binding Protein. mBio. 2020;11(3):10.1128/mbio.00810-20.

63. Brocca S, Grandori R, Longhi S, Uversky V. Liquid–Liquid Phase Separation by Intrinsically Disordered Protein Regions of Viruses: Roles in Viral Life Cycle and Control of Virus–Host Interactions. International Journal of Molecular Sciences [Internet]. 2020; 21(23).

64. Tollervey D, Lehtonen H, Jansen R, Kern H, Hurt EC. Temperature-sensitive mutations demonstrate roles for yeast fibrillarin in pre-rRNA processing, pre-rRNA methylation, and ribosome assembly. Cell. 1993;72(3):443–57.

65. Tessarz P, Santos-Rosa H, Robson SC, Sylvestersen KB, Nelson CJ, Nielsen ML, et al. Glutamine methylation in histone H2A is an RNA-polymerase-I-dedicated modification. Nature. 2014;505(7484):564-8.

66. Banerjee J, Radvar E, Azevedo HS. Self-assembling peptides and their application in tissue engineering and regenerative medicine. In: Barbosa MA, Martins CL, editors. Peptides and Proteins as Biomaterials for Tissue Regenaration and Repair: WoodHead Publishing, Elsevier; 2018. p. 245–81.

67. Tanford C. Contribution of Hydrophobic Interactions to the Stability of the Globular Conformation of Proteins. Journal of the American Chemical Society. 1962;84(22):4240–7.

68. Bresnick EH, Felsenfeld G. The leucine zipper is necessary for stabilizing a dimer of the helix-loop-helix transcription factor USF but not for maintenance of an elongated conformation. J Biol Chem. 1994;269(33):21110–6.

69. Situ AJ, Kang SM, Frey BB, An W, Kim C, Ulmer TS. Membrane Anchoring of α-Helical Proteins: Role of Tryptophan. J Phys Chem B. 2018;122(3):1185–94.

70. Zhang Q, Huang H, Zhang L, Wu R, Chung CI, Zhang SQ, et al. Visualizing Dynamics of Cell Signaling In Vivo with a Phase Separation-Based Kinase Reporter. Mol Cell. 2018;69(2):347.

71. Maurer AC, Cepeda Diaz AK, Vandenberghe LH. Residues on Adeno-associated Virus Capsid Lumen Dictate Interactions and Compatibility with the Assembly-Activating Protein. Journal of Virology. 2019;93(7):1–16.

72. Vogel R, Seyffert M, Strasser R, de Oliveira Anna P, Dresch C, Glauser Daniel L, et al. Adeno-Associated Virus Type 2 Modulates the Host DNA Damage Response Induced by Herpes Simplex Virus 1 during Coinfection. Journal of Virology. 2012;86(1):143–55.

73. Zolotukhin S, Byrne BJ, Mason E, Zolotukhin I, Potter M, Chesnut K, et al. Recombinant adeno-associated virus purification using novel methods improves infectious titer and yield. Gene Therapy. 1999;6(6):973–85.

74. Laughlin CA, Tratschin J-D, Coon H, Carter BJ. Cloning of infectious adeno-associated virus genomes in bacterial plasmids. Gene. 1983;23(1):65–73.

75. Sutter SO, Marconi P, Meier AF. Herpes Simplex Virus Growth, Preparation, and Assay. In: Diefenbach RJ, Fraefel C, editors. Herpes Simplex Virus : Methods and Protocols. New York, NY: Springer New York; 2020. p. 57–72.

76. Laughlin CA, Tratschin JD, Coon H, Carter BJ. Cloning of infectious adeno-associated virus genomes in bacterial plasmids. Gene. 1983;23(1):65–73.

77. Schindelin J, Arganda-Carreras I, Frise E, Kaynig V, Longair M, Pietzsch T, et al. Fiji: an open-source platform for biological-image analysis. Nature Methods. 2012;9(7):676–82.

78. Vetter J, Papa G, Tobler K, Rodriguez Javier M, Kley M, Myers M, et al. The recruitment of TRiC chaperonin in rotavirus viroplasms correlates with virus replication. mBio. 2024;0(0):e00499–24.

79. Koulouras G, Panagopoulos A, Rapsomaniki MA, Giakoumakis NN, Taraviras S, Lygerou Z. EasyFRAP-web: a web-based tool for the analysis of fluorescence recovery after photobleaching data. Nucleic Acids Research. 2018;46(W1):W467–W72.

80. Eichwald C, Kim J, Nibert ML. Dissection of mammalian orthoreovirus µ2 reveals a self-associative domain required for binding to microtubules but not to factory matrix protein µNS. PLoS ONE. 2017;12(9):1–25.

81. Mirdita M, Schütze K, Moriwaki Y, Heo L, Ovchinnikov S, Steinegger M. ColabFold: making protein folding accessible to all. Nature Methods 2022 19:6. 2022;19(6):679–82.

