## Supplementary material for "Assembly-Activating Protein Phase Separation Properties Are Required for Adeno-Associated Virus Type 2 Assembly": suppl Fig S1, S2,S3, S4 and corresponding Fig Legends and Tables S1 and S2

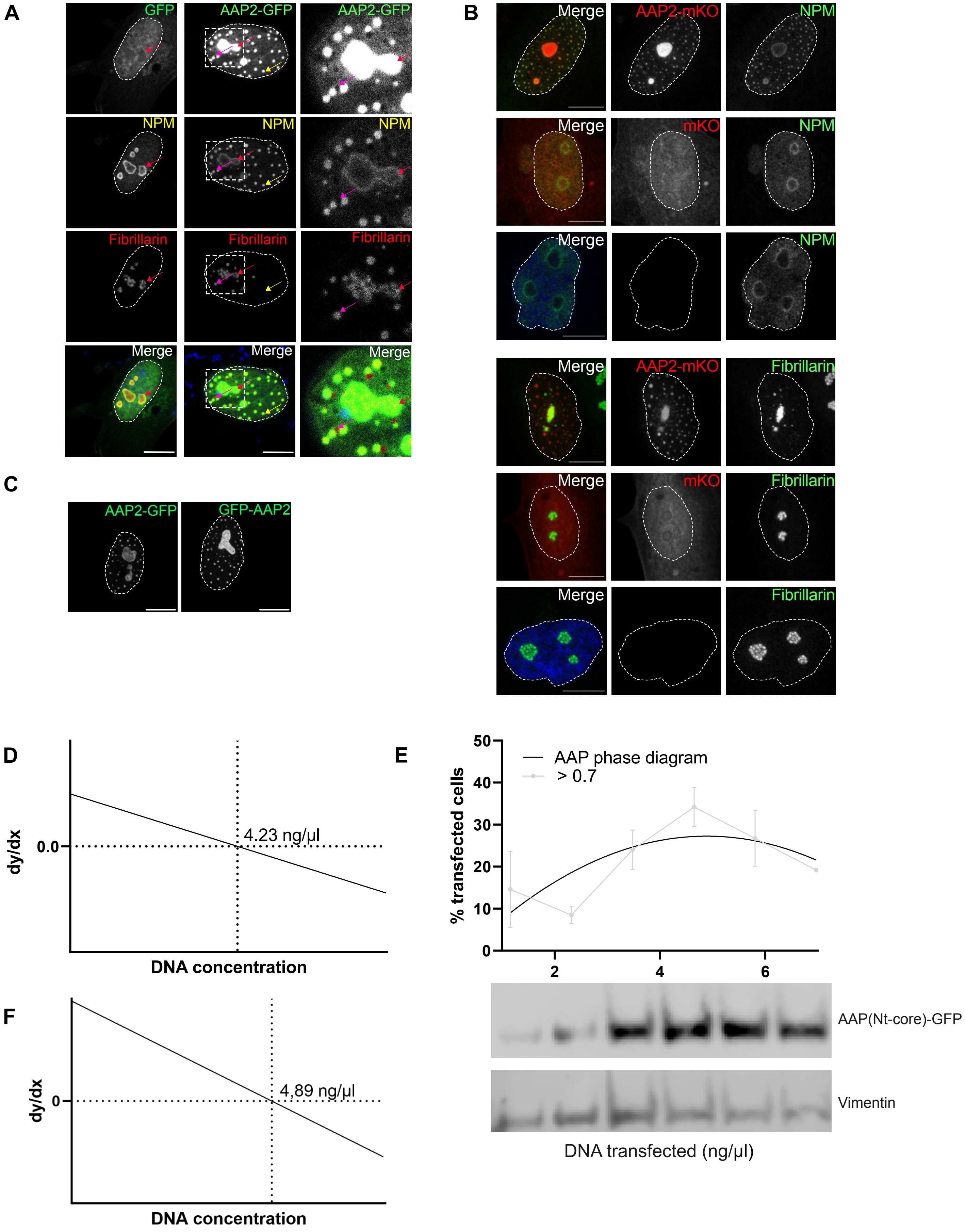

A
GFP
AAP2-GFP
AAP2-GFP
NPM
NPM
NPM
Fibrillarin
Fibrillarin
Fibrillarin
Merge
Merge
Merge
B
Merge
AAP2-mKO
NPM
Merge
mKO
NPM
Merge
NPM
Merge
AAP2-mKO
Fibrillarin
Merge
mKO
Fibrillarin
Merge
Fibrillarin
C
AAP2-GFP
GFP-AAP2
D
dy/dx
0.0
4.23 ng/µl
DNA concentration
E
% transfected cells
AAP phase diagram
> 0.7
50
40
30
20
10
0
2
4
6
AAP(Nt-core)-GFP
Vimentin
DNA transfected (ng/µl)
F
dy/dx
0
4.89 ng/µl
DNA concentration

Fig S2

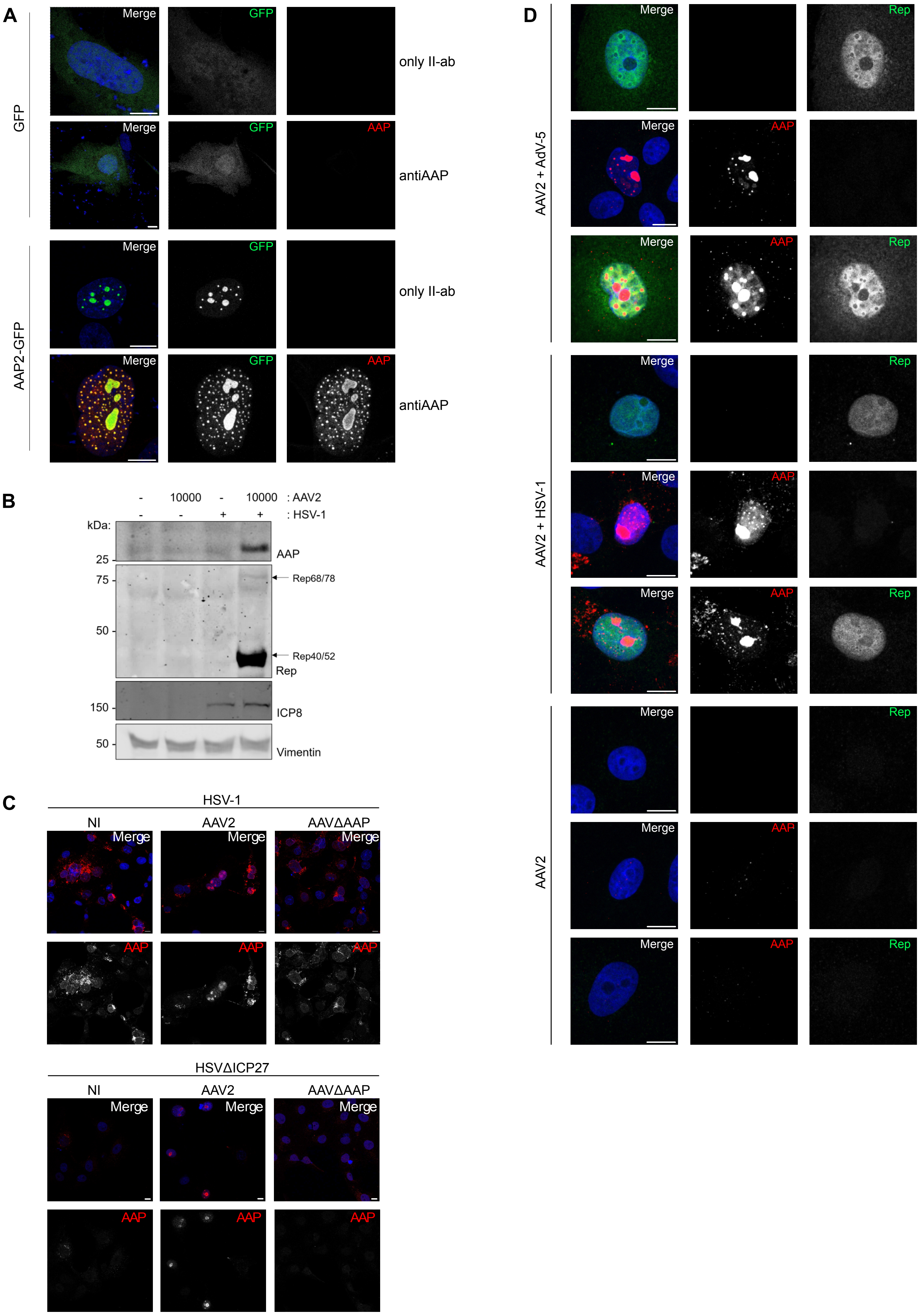
A
GFP
AAP2-GFP
Merge
GFP
AAP
only II-ab
antiAAP
B
- 10000 - 10000 : AAV2
- - + + : HSV-1
kDa:
25
75
50
150
50
AAP
Rep68/78
Rep40/52
Rep
ICP8
Vimentin
C
HSV-1
NI
AAV2
AAVΔAAP
HSVΔICP27
D
AAV2 + AdV-5
AAV2 + HSV-1
AAV2
Rep

### Figure S3

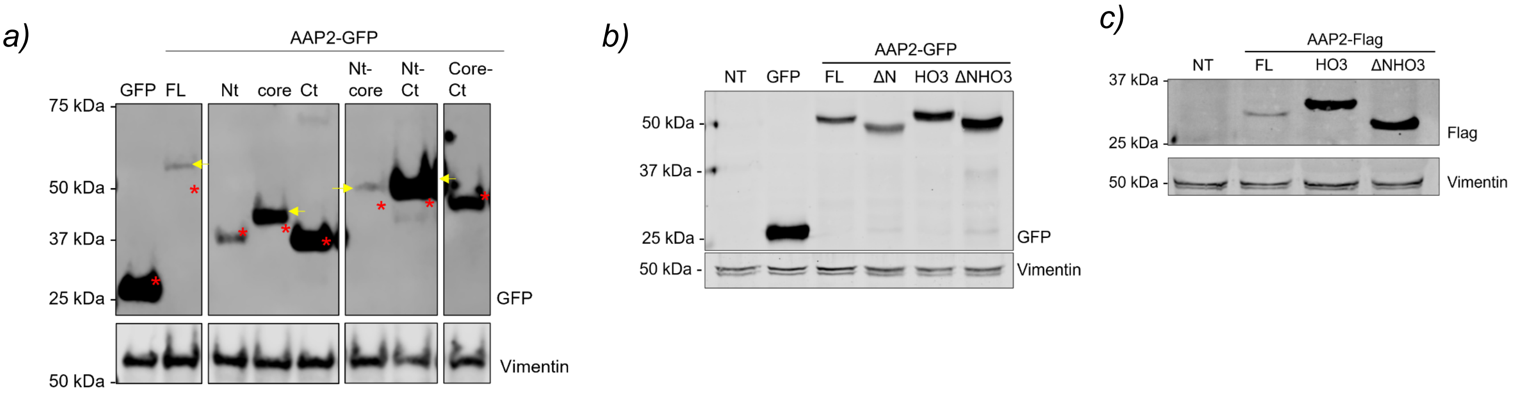


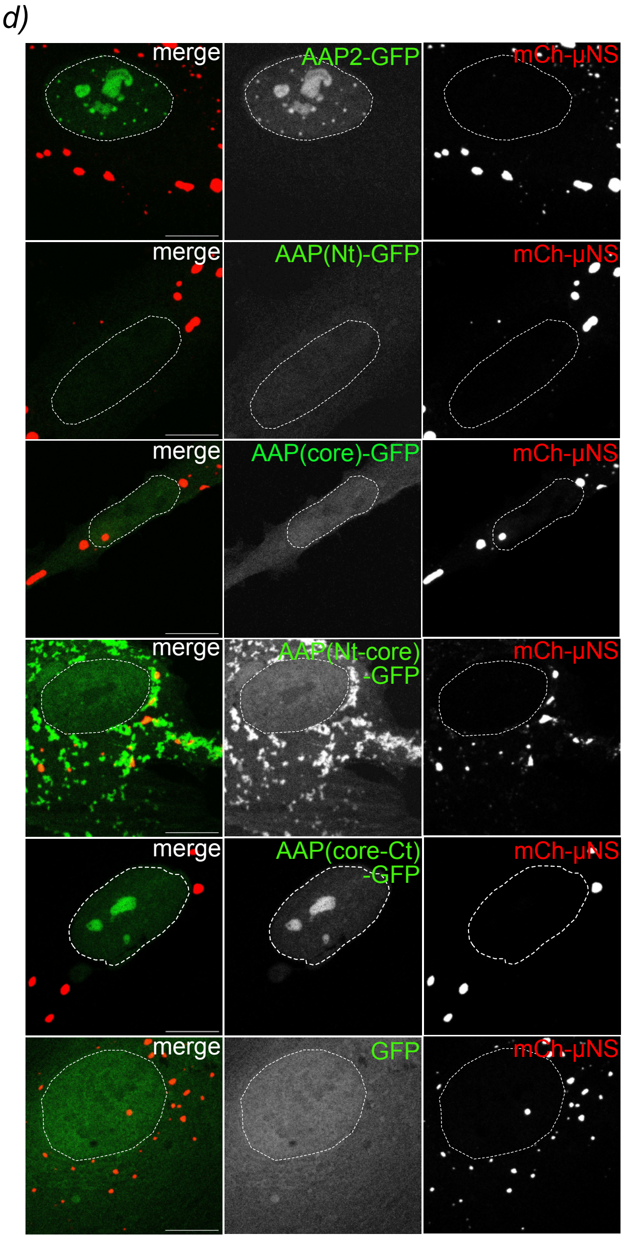


e)

| Protein | Predicted size (kDa) |
|---|---|
| GFP | 27 |
| AAP-GFP | 50 |
| AAP(Nt)-GFP | 36 |
| AAP(core)-GFP | 36 |
| AAP(Ct)-GFP | 34 |
| AAP(Nt-core)-GFP | 43 |
| AAP(Nt-Ct)-GFP | 42 |
| AAP(core-Ct)-GFP | 45 |
| AAP(PM)-GFP | 50 |
| ΔNAAP-GFP | 48 |
| AAP(HO3)-GFP | 51 |
| ΔNAAP(HO3)-GFP | 48 |
| AAP-Flag | 24 |
| AAP(HO3)-Flag | 25 |
| ΔNAAP(HO3)-Flag | 23 |

Fig S4

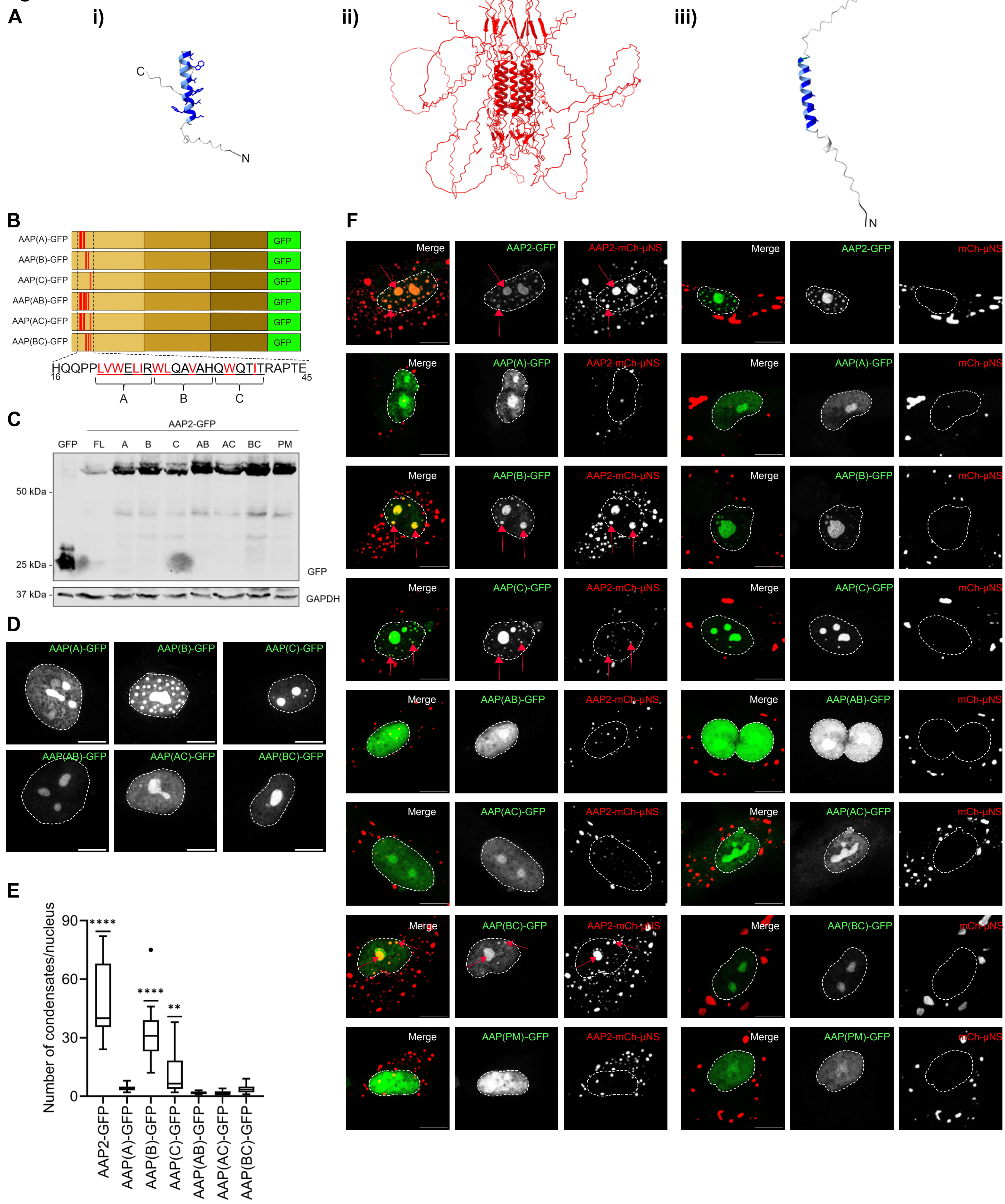

A
i)
ii)
iii)
C
N
B
AAP(A)-GFP
AAP(B)-GFP
AAP(C)-GFP
AAP(AB)-GFP
AAP(AC)-GFP
AAP(BC)-GFP
GFP
HQQPPLVWELIRWLQAVAHQWQTITRAPTE
16
45
A
B
C
C
AAP2-GFP
GFP FL A B C AB AC BC PM
50 kDa
25 kDa
37 kDa
GFP
GAPDH
D
E
Number of condensates/nucleus
90
60
30
0
****
****
**
AAP2-GFP
AAP(A)-GFP
AAP(B)-GFP
AAP(C)-GFP
AAP(AB)-GFP
AAP(AC)-GFP
AAP(BC)-GFP
F
Merge
AAP2-GFP
AAP2-mCh-μNS
mCh-μNS
AAP(A)-GFP
AAP(B)-GFP
AAP(C)-GFP
AAP(AB)-GFP
AAP(AC)-GFP
AAP(BC)-GFP
AAP(PM)-GFP

**Supporting Information**

**Supporting Figure Legends**

**Fig S1. AAP2 inclusions and phase diagrams.** (A) Immunofluorescence images of BJ cells expressing GFP (green, left column) or AAP2-GFP (green, right column). At 16 hpt, the cells were fixed and immunostained for the detection of nucleolar proteins: nucleophosmin (anti-NPM, yellow) and fibrillarin (anti-fibrillarin, red). The nuclei were stained with DAPI (blue). A merged image is shown at the bottom. The dashed open box corresponds to the enlarged picture from the right column. Red, yellow, and pink arrows point to nucleoli, extranucleolar inclusions, and dispersed fibrillarin, respectively. The scale bar is 10 µm. (B) Immunofluorescence images of BJ cells expressing AAP2-mKO (red, top row), mKO (red, middle row), or of untransfected cells (bottom row). At 16 hpt, the cells were fixed and immunostained for the detection of nucleolar proteins: nucleophosmin (anti-NPM, green, top panel) or fibrillarin (anti-fibrillarin, green, bottom panel). Nuclei were stained with DAPI (blue, dashed white line). A merged image is shown in the left column. The scale bar is 10 µm. (C) Immunofluorescence images of BJ cells expressing AAP2-GFP (left image) or GFP-AAP2 (right image). At 16 hpt, the cells were fixed, and nuclei were stained with DAPI (dashed white line). The scale bar is 10 µm. (D) Plot of the first derivative of the AAP2-GFP phase diagram, having a maximal phase separation with a DNA concentration of 4.23 ng/µl. (E) Plot of the phase diagram of AAP(Nt)-GFP obtained from BJ cells transfected with pCI-AAP(Nt-core)-GFP at increasing DNA concentrations. The AAP(Nt-core)-GFP condensate fractions had a cut-off with a solidity over 0.7 (grey line). The data corresponds to three independent experiments. At the bottom, immunoblotting for the expression of AAP(Nt-core)-GFP (anti-GFP). Vimentin was used as a loading control. (F) Plot of the first derivative of the AAP(Nt-core)-GFP phase diagram, having a maximal phase separation with a DNA concentration of 4.89 ng/µl.

**Fig S2. Anti-AAP antibody can recognize AAP2 in immunofluorescence and immunoblotting.** (A) Immunofluorescence images of BJ cells expressing GFP (top panel) or AAP2-GFP (bottom panel). At 16 hpt, the cells were paraformaldehyde-fixed and immunostained for the detection of GFP and AAP2 with anti-AAP (second and fourth rows). The GFP signal is green). The nuclei were stained with DAPI (blue). The scale bar is 10 μm. (B) Immunoblotting of Vero cell extracts at 24 hpi with AAV2 (MOI 10000) in absence or presence of HSV-1 (MOI 0.1). The membrane was incubated with anti-AAP, anti-Rep, and anti-ICP8. Anti-vimentin was used as a loading control. (C) Immunofluorescence images of Vero cells infected with HSV-1 (MOI 0.1, top panel) or HSVΔICP27 (MOI 1, bottom panel) alone or co-infected with HSV-1 (MOI 0.1, top panel) or HSVΔICP27 (MOI 1, bottom panel) and either AAV2 (MOI 500) or AAVΔAAP (MOI 500). At 24 hpi, the cells were paraformaldehyde-fixed and immunostained for the detection of AAP2 (anti-AAP, red). The nuclei were stained with DAPI (blue). The scale bar is 10 μm. (D) Maximal projection of immunofluorescence images from Z-stacks of Vero cells co-infected with AAV2 (MOI 500) and either AdV-5 (MOI 0.1, top panel) or HSV-1 (MOI 0.1, middle panel) or infected with AAV2 alone (MOI 500, bottom panel). At 24 hpi, the cells were paraformaldehyde-fixed and immunostained for the detection of Rep proteins (anti-Rep, green, top, and middle rows) and AAP2 (anti-AAP, red, middle, and bottom rows). The nuclei were stained with DAPI (blue). A merge image is shown in the left column. The scale bar is 10 μm.

**Fig S3. Characterization of the expression of AAP2 deletion mutants.** (A) Immunoblotting of extracts from BJ cells expressing AAP2-GFP or deletions mutants of AAP2-GFP. The membrane was incubated with anti-GFP (top panel) and, as loading control, anti-vimentin (bottom panel). The red stars point to the predicted molecular weight of the corresponding protein. The red arrow indicates a slow migration form of AAP2-GFP, probably due to a post-translational modification. (B) Immunoblotting of extracts from Vero cells expressing APP2-GFP, ΔNAAP-GFP, AAP(HO3)-GFP, or ΔNAAP(HO3)-GFP. The membrane was incubated with anti-GFP (top panel) and anti-vimentin (bottom panel) as loading control. The red stars point to the predicted molecular weight of the corresponding protein. (C) Immunoblotting of extracts from Vero cells expressing AAP2-Flag, AAP(HO3)-Flag, or ΔNAAP(HO3)-Flag. The membrane was incubated with anti-Flag (top panel) and, as loading control, anti-vimentin (bottom panel). The red stars point to the predicted molecular weight of the corresponding protein. (D) Immunofluorescence images of control AAP2-GFP or AAP2-GFP deletion mutants with mCherry-µNS in the reovirus platform. For this, BJ cells co-expressing AAP2-GFP or AAP2-GFP deletion mutants (green) with mCherry-µNS (red) were fixed, followed by staining of the nuclei with DAPI (white dashed line). A merged image is shown in the left column. The scale bar is 10 µm. (E) Table indicating the predicted molecular weight of AAP2-GFP and diverse AAP2-GFP deletion mutants.

**Fig S4. AAP2 point mutants are deficient in BC formation.** (A) AlphaFold2 predictions of i) the monomeric AAP N-terminus (amino acid region 1 to 60), bulky hydrophobic residues are indicated in dark blue; ii) the oligomeric AAP2 as pentamer, each monomeric AAP is in red; and iii) the monomeric AAP (PM) N-terminus (amino acid region 1 to 60), having substituted in predicted α-helix the hydrophobic bulky residues by alanines. The values of the predicted local distance difference test (pLDDT) are shown as color on each predicted structure: blue (pLDDT >50) and red (pLDDT <50). (B) Schematic representation of AAP2-GFP with hydrophobic residues substituted by alanines in groups A, B, or C and their combinations. The substituted hydrophobic residues are indicated in red. (C) Immunoblotting of extract from BJ cells expressing indicated AAP2-GFP with hydrophobic residues substituted by alanines. The membrane was incubated for the detection of AAP2-GFP (anti-GFP, top panel) and, as loading control, GAPDH (anti-GAPDH, bottom panel). (D) Immunofluorescence images of BJ cells expressing AAP2-GFP with indicated point mutations. At 16 hpt, the cells were fixed, and nuclei were stained with DAPI (dashed white line). The scale bar is 10 µm. (E) Plot of the numbers of AAP2-GFP condensates harboring the indicated point mutations. The data represent the mean ± SD; one-way ANOVA, (**) $p < 0.01$, (****) $p < 0.0001$, $n \geq 15$. (F) Immunofluorescence images of *in vivo* protein interaction using reovirus platform between indicated AAP2-GFP point mutations (green) with AAP2-mCherry-µNS (red, left panel) or mCherry-µNS (red, right panel). At 16 hpt, the BJ cells were fixed, and nuclei were stained with DAPI (dashed white line). A merged image is shown on the left side of each column. The red arrows point to colocalization of AAP2-GFP point mutations with the AAP2-mCherry-μNS platform. The scale bar is 10 μm.

#### Supporting Tables

**Table S1.** Oligonucleotide sequences used in this study.

| Segment | Oligonucleotide sequence |
|---|---|
| AAP2 | fwd: 5'-gatcgcctcgaggccacc**atg**gagacgcagactcagtacctg-3' |
| | rev: 5'-gatcacgcgtgggtgaggtatccatactgtg-3' |
| GFP | fwd: 5'-agctacgcgt**atg**gtgagcaagggcgaggag-3' |
| | rev: 5'-gatcgcggccgc**tta**cttgtacagctcgtccatg-3' |
| mKO2 | fwd: 5'-gatctctaga**atg**gtgagtgtgattaaacca-3' |
| | rev: 5'-gatcgcggccgc**tta**atgagctactgcatcttctac-3' |
| AAP(Nt) | fwd: 5'-tataggctagcgccgcc**atg**gagacgcagactcagtac-3' |
| | rev: 5'-accatacgcgtcatcgtgtcgacccatccatgtggaatcgcaatg-3' |
| AAP(core) | fwd: 5'-tataggctagcgccgcc**atg**gcgacagagtcatcaccacc-3' |
| | rev: 5'-accatacgcgtgaatgttaaagagcttgaagttgagtctc-3' |
| AAP(Ct) | fwd: 5'-tataggctagcgtcgacgccgcc**atg**aagtcaaagaggtcacgcag-3' |
| | rev: 5'-accatacgcgtgggtgaggtatccatactgtgg-3' |
| 6xHis tag | fwd: 5'-cgcgtcgtggcagccatcaccatcaccatcac**tga**gc-3' |
| | rev: 5'-ggccgc**tca**gtgatggtgatggtgatggctgccacga-3' |
| Flag tag | fwd: 5'-cgcgtgactataaggacgatgatgacaaa**taa**gc-3' |
| | rev: 5'-ggccgc**tta**tttgtcatcatcgtccttatagtca-3' |
| Stop codon | fwd: 5'-cgcg**ttaataa**gc-3' |
| | rev: 5'-ggccgc**ttatta**a-3' |

* Restriction enzyme sites are underlined.

** Initiation and stop codons are labeled in bold.

**Table S2.** Synthetic gene fragments used in this study.

| Name of gene segment | Gene fragment sequence |
|---|---|
| ΔNAAP | 5’-ataggctagcctcgaggccaccatgctggtctgggaactaatacgatggc<br>tacaggcagtggcgcaccaatggcagacaataacgagggcgccgacggagtgggtaattcctcg<br>ggaaatt-3’ |
| AAP(HO3) | 5’-ataggctagcctcgaggccaccatggagacgcagactcagtacctgaccc<br>ccagcctctcggacagccaccagcagccccctggagaaatcgcaaagagtctgaaagaaattgc<br>aaaatccttgaaggaaatagcgtggtcactgaaagaaattgccaagtccctcaaagggagggcg<br>ccgacggagtgggtaattcctcgggaaatt-3’ |
| ΔNAAP(HO3) | 5’-ataggctagcctcgaggccaccatgggagaaatcgcaaagagtctgaaag<br>aaattgcaaaatccttgaaggaaatagcgtggtcactgaaagaaattgccaagtccctcaaaggg<br>agggcgccgacggagtgggtaattcctcgggaaatt-3’ |
| AAP(PM) | 5’-ataggctagcgccgccatggagacgcagactcagtacctgacccccagc<br>ctctcggacagccaccagcagccccctgcagccgcagaagcagcacgagctgcacaggcagc<br>ggcgcaccaagcacagacagcaacgagggcgccgacggagtgggtaattcctcgggaaatt-3’ |
| AAP(A) | 5’-ataggctagcgccgccatggagacgcagactcagtacctgacccccagc<br>ctctcggacagccaccagcagccccctgcagccgcagaagcagcacgatggctacaggcagtg<br>gcgcaccaatggcagacaataacgagggcgccgacggagtgggtaattcctcgggaaatt-3’ |
| AAP(B) | 5’-ataggctagcgccgccatggagacgcagactcagtacctgacccccagcc<br>tctcggacagccaccagcagccccctctggtctgggaactaatacgagctgcacaggcagcggc<br>gcaccaatggcagacaataacgagggcgccgacggagtgggtaattcctcgggaaatt-3’ |
| AAP(C) | 5’-ataggctagcgccgccatggagacgcagactcagtacctgacccccagcc<br>tctcggacagccaccagcagccccctctggtctgggaactaatacgatggctacaggcagtggcg<br>caccaagcacagacagcaacgagggcgccgacggaggcagcaattcctcgggaaatt-3’ |

| | |
|---|---|
| AAP(AB) | 5’-ataggctagcgccgccatggagacgcagactcagtacctgacccccagcc<br>tctcggacagccaccagcagccccctgcagccgcagaagcagcacgagctgcacaggcagcg<br>gcgcaccaatggcagacaataacgagggcgccgacggagtgggtaattcctcgggaaatt-3’ |
| AAP(AC) | 5’-ataggctagcgccgccatggagacgcagactcagtacctgacccccagcc<br>tctcggacagccaccagcagccccctgcagccgcagaagcagcacgatggctacaggcagtgg<br>cgcaccaagcacagacagcaacgagggcgccgacggaggcagcaattcctcgggaaatt-3’ |
| AAP(BC) | 5’-ataggctagcgccgccatggagacgcagactcagtacctgacccccagcc<br>tctcggacagccaccagcagccccctctggtctgggaactaatacgagctgcacaggcagcggc<br>gcaccaagcacagacagcaacgagggcgccgacggaggcagcaattcctcgggaaatt-3’ |

* The sequence corresponds to the upper strand of each synthetic gene fragment.
